# Activation of M_1_ muscarinic receptors reduce pathology and slow progression of neurodegenerative disease

**DOI:** 10.1101/2021.07.30.454298

**Authors:** Louis Dwomoh, Mario Rossi, Miriam Scarpa, Elham Khajehali, Colin Molloy, Pawel Herzyk, Shailesh N. Mistry, Andrew R. Bottrill, Patrick M. Sexton, Arthur Christopoulos, P. Jeffery Conn, Craig W. Lindsley, Sophie J. Bradley, Andrew B. Tobin

## Abstract

The most prevalent types of dementias, including Alzheimer’s disease, are those that are propagated via the spread of *“prion-like”* misfolded proteins. Despite considerable effort no treatments are available to slow or stop the progression of these dementias. Here we investigate the possibility that activation of the M_1_-muscarinic receptor (M_1_-receptor), which is highly expressed in the brain and that shows pro-cognitive properties, might present a novel disease modifying target. We demonstrate that the progression of murine prion disease, which we show here displays many of the pathological, behavioural and biochemical hallmarks of human neurodegenerative disease, is slowed and normal behaviour maintained by the activation of the M_1_-receptor with a highly tolerated positive allosteric modulator (VU846). This correlates with a reduction in both neuroinflammation and indicators of mitochondrial dysregulation, as well as a normalisation in the expression of markers associated with neurodegeneration and Alzheimer’s disease. Furthermore, VU846 preserves expression of synaptic proteins and post-synaptic signalling components that are altered in disease. We conclude that allosteric regulation of M_1_-receptors has the potential to reduce the severity of neurodegenerative diseases caused by the *prion-like* propagation of misfolded protein in a manner that extends life span and maintains normal behaviour.

## INTRODUCTION

Alzheimer’s disease (AD) is the most prevalent form of neurodegenerative disease and is predicted to double every 20 years in line with an aging global population ^1^. Despite the progress made in our understanding of AD genetic risk factors ^2–4^ and in the identification of disease-associated systems ^5^ there are currently no treatments that can stop or slow the progression of AD ^6^. Among the most prevalent of the adaptive mechanisms in operation in neurodegenerative disorders are neuroinflammation ^7^ and the surveillance and clearance of misfolded protein ^8^. Although there is a growing appreciation of the key cellular and protein components underpinning these processes, designing effective clinical strategies that target these processes in a manner that will slow or halt neurodegenerative disease remains a major challenge.

The key pathological characteristic of AD is a loss of acetylcholine-cholinergic neurons originating from basal forebrain nuclei innervating limbic and neocortical structures ^9,10^. Current frontline treatments are focused on overcoming the ensuing deficit in cholinergic transmission by elevating brain acetylcholine levels through the inhibition of cholinesterases, the enzymes responsible for acetylcholine catabolism ^2^. Whereas this approach has some limited clinical benefit in the symptomatic treatment of memory loss in early stages of disease ^11–13^, there is no consistent evidence that this approach can slow the progression of disease. Despite this there are emerging reports that activation of post-synaptic acetylcholine receptors of the muscarinic receptor family, which consist of five receptor subtypes (M_1_-M_5_-receptors), but particularly the M_1_-receptor subtype, can offer neuroprotection in the context of neurodegenerative disease ^14, 15^. This is particularly exciting given that activation of the M_1_-receptor is widely considered as a promising strategy for the treatment of memory loss in AD due to high expression of this receptor subtype in the cortex and hippocampus ^14, 16^, and robust pro-cognitive effects following receptor activation in pre-clinical animal models ^14, 17–19^. Combined, these studies suggest that selective targeting of the M_1_-receptor might have a dual benefit in AD through restoration of cognitive function, and neuroprotection that would slow disease progression.

The barrier to testing this hypothesis in the clinic is the development of agonists that selectively activate the M_1_-receptor since the orthosteric acetylcholine binding site is nearly identical between the 5 muscarinic receptor subtypes ^20^. As such, generation of subtype-selective orthosteric agonists is very challenging ^21^. An alternative approach we, and others, have adopted is to target an allosteric binding pocket on the extracellular surface of the receptor ^20^. Agents that bind to this site can act by increasing the sensitivity of the receptor to acetylcholine ^22^. These positive allosteric modulators (PAMs) have the advantage of being highly selective for the M_1_-receptor while maintaining the spatiotemporal profile of cholinergic signalling since they act co-operatively with the natural ligand acetylcholine. These features are considered to be the primary reasons for PAMs showing reduced adverse responses normally associated with prolonged activation by orthosteric M_1_-receptor agonists ^23, 24^.

Our initial studies investigated the activity the prototypical M_1_-receptor PAM, benzyl quinolone carboxylic acid (BQCA) ^19^, in murine prion disease which is a terminal neurodegenerative disease where there is a progressive loss of hippocampal neurons, ^25^ including a disruption of hippocampal-cholinergic innervation with associated deficits in learning and memory, that we have shown can be restored by treatment with clinical cholinesterase inhibitors ^14^. In this model we found that a single-dose of BQCA prior to training in a fear conditioning protocol restored defective learning and memory in murine prion disease whilst chronic daily treatment for several weeks slowed disease progression ^14^.

Here we substantially extend these earlier studies using global proteomic and transcriptomic analysis of murine prion disease to report that this model shows adaptive-responses, including neuroinflammation and up-regulation of protein markers of AD, as well as indicators of synaptic loss and mitochondrial dysfunction, that are characteristic hallmarks of AD. We further report that the next generation M_1_-PAM, VU0486846 (VU846) ^26, 27^, reduced markers of neuroinflammation and prevented the up-regulation of adaptive disease responses whilst maintaining synaptic proteins at near normal levels. These observations offered mechanistic insight into the improvement in behavioural symptoms and the extension of survival of prion-diseased mice associated with administration of M_1_-receptor PAMs. We conclude that M_1_-PAMs exhibit therapeutic potential for slowing the progression of neurodegenerative disease by regulating adaptive responses such as neuroinflammation that are common features of the pathology of brain diseases caused by the propagation of misfolded protein.

## RESULTS

### M_1_-PAMs restore learning and memory and prolong survival in murine prion disease

We have previously reported that defective learning and memory associated with a disruption of hippocampal cholinergic innervation in murine prion disease can be restored by orthosteric and allosteric M_1_-receptor ligands ^14^. This was observed in Tg37 mice, an engineered strain that over-expresses (3x) murine prion protein, infected with prion-brain homogenate (rocky mountain laboratory (RML)) or control normal brain homogenate (NBH) ^25^. In this system the M_1_-receptor selective PAM, BQCA, was shown to be highly tolerated and following chronic (daily) dosing, prolonged survival of prion diseased mice ^14^. Here we extend these studies by using a next generation M_1_-PAM, VU846 ^27^, that in cortical neuronal cultures showed high co-operativity with acetylcholine (Log*αβ*=1.38, n=4) in second messenger, IP1, assays **(Fig 1A,B)**. In addition, a single administration of VU846 (10mg/kg) 30 minutes prior to fear-conditioning training completely restored defective contextual fear conditioning learning and memory in prion diseased animals **(Fig 1C)**.

**Figure 1.**
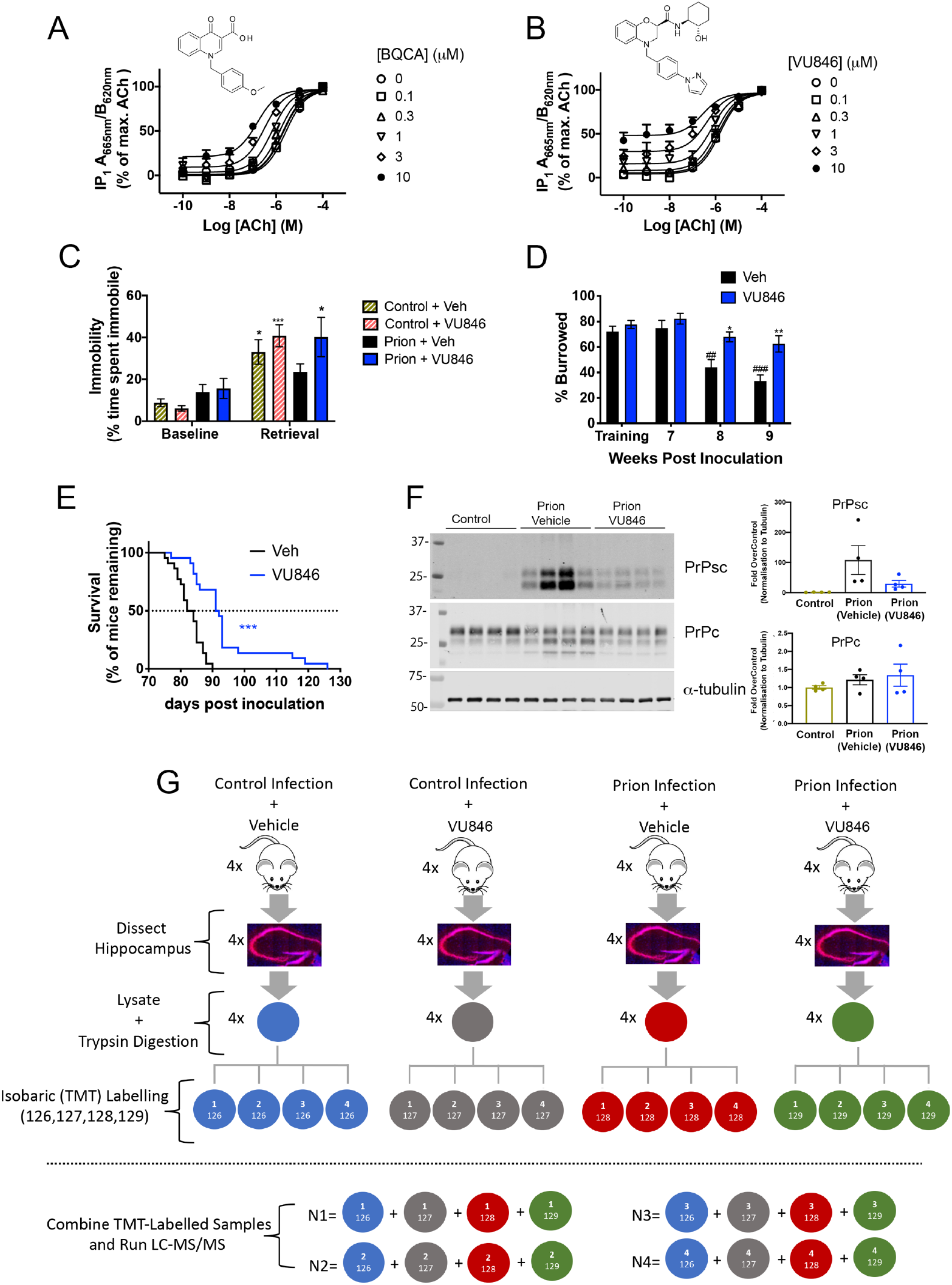
M1-receptor PAMs restore learning and memory and prolong survival in murine prion disease. M1-receptor PAMs, (**A**) BQCA and (**B**) VU846 enhance the ACh-mediated IP1 accumulation in cortical neuronal cultures. Data shown are the mean + SEM. n=4. (**C**) Fear-conditioning response of control and prion-infected mice following acute administration of vehicle (20% tween-80) or VU846 (10 mg/kg) prior to training and retrieval. n=15-19 mice per group. 2-way ANOVA with Sidak’s multiple comparison test. \*\*\**P*<0.001, \**P*<0.05 vs. control + vehicle. Data are shown as mean ± SEM. (**D**) Burrowing response of control and prion-infected mice following administration of vehicle or VU846 (10 mg/kg) 30 minutes before each burrowing session (from 7 w.p.i.). n=12-13 mice per group. ##*P*<0.01; ###*P*<0.001 w.p.i. vs. training, \**P*<0.05; \*\**P*<0.01 prion-vehicle vs. prion-VU846. (**E**) Kaplan-Meier survival plots for prion-infected mice treated with vehicle (20% tween-80) n = 22; black line) or VU846 (10 mg/kg; n = 22; blue line) daily from 7 w.p.i. \*\*\**P* < 0.001, Gehan-Breslow-Wilcoxon test. (**F**) Lysates from control, prion-vehicle and prion-VU846 (10 mg/kg) mouse hippocampus were probed in western blots with an antibody that detected both cellular (PrP_c_) and misfolded proteinase K–resistant scrapie (PrP_sc_) prion protein. The upper gel displays lysates treated with proteinase K before probing in western blots for prion protein. The lower gel displays the same lysates but not treated with proteinase K. Each lane is from a different animal. Quantitative data from the blots are shown as bar graphs of mean ± SEM, with individual values also displayed. n=4. (**G**) Illustrated summary of the experimental outline and sample preparation for TMT mass spectrometry-based proteomics.

We next tested if VU846 had the potential to modify the progression of prion disease. In these experiments animals were dosed daily with VU846 (10mg/kg) from 7 weeks post inoculation (w.p.i.), a time point where animals showed the first pathological signs of misfolded scrappie prion protein (PrP_sc_) **(Supplementary Fig S1A-C)** but no physical indicators of disease. Animals treated with vehicle showed reduced performance in burrowing, an innate behaviour associated with hippocampal function, whilst in prion infected animals treated with VU846 burrowing behaviour was significantly improved when compared with vehicle treated controls **(Fig 1D)**. Furthermore, there was a significant delay in the onset of terminal clinical symptoms in animals dosed daily with VU846 **(Fig 1E)** – with some animals showing very extended life spans **(Fig 1E)**. A slowing in disease progression was further evident in the observed reduction in accumulation of PrP_sc_ in VU846 treated animals **(Fig 1F).**

In summary, these studies established that the M_1_-PAM, VU846, restored learning and memory in murine prion disease when administered acutely and possessed disease modifying properties that corrected for behaviour abnormalities and promoted survival on chronic exposure.

### Markers of neurodegeneration and neuroinflammation that overlap those in Alzheimer’s disease are up-regulated in murine prion disease

We next determined changes to the proteome caused by prion infection; called here the *prion-effect.* This was conducted on hippocampi isolated from Tg37 mice inoculated with RML or as a control, NBH, at 3 weeks of age and then treated with vehicle (i.p.) from 7-10 w.p.i **(Fig 1G)**. Principal component analysis showed good separation of the control animals from the prion infected animals **(Fig 2A)**. The total number of proteins identified was 6,208 of which 4,528 met our robustness criteria and were quantified **(Supplementary Table S1).** Of these, 566 proteins were significantly up-regulated by more than 0.4 log2 fold in prion disease whilst 10 proteins were down-regulated (p<0.05 (-log_10_ 0.05 = 1.30)) **(Fig 2B, Supplementary Table S2).** Go-ontology (GO) analysis showed that the proteins up-regulated in prion disease fell into enrichment groups associated with processes known to be up-regulated in human neurodegenerative diseases **(Fig 2C, Supplementary Table S3).** These include groups associated with neuronal death, synaptic pruning, neurofibrillary tangle assembly, oxidative stress and protease activity **(Fig 2C).** In addition, markers of neuroinflammation and in particular indicators of astrocyte and microglia activation (e.g. vimentin, galectin-1 and glial fibrillary acidic protein (GFAP)) were up-regulated in murine prion disease **(Fig 2B,C)**. The activation of neuroinflammatory pathways was also evident in the up-regulation of components of the neuronal complement system, (e.g. C1qA,B,C) **(Fig 2B,C)**. Significantly, many of these neuroinflammatory markers are also reported to be up-regulated in AD ^28, 29^. Furthermore, the *prion-effect* included an up-regulation of other proteins considered markers of AD ^28, 29^. This included, for example, the up-regulation of several members of the apolipoprotein family **(Fig 2B, Supplementary Table S2 and S3)** including ApoE. This is consistent with reports from human prion disease where ApoE was also reported to be up-regulated ^30, 31^. Furthermore, key enzymes and transporters such as transglutaminase 1 (Tgm1) and ABCA1, that have been implicated in the clearance of misfolded protein in AD ^32, 33^, are also seen to be up-regulated in murine prion disease **(Fig 2B, Supplementary Table S2 and S3)**.

**Figure 2.**
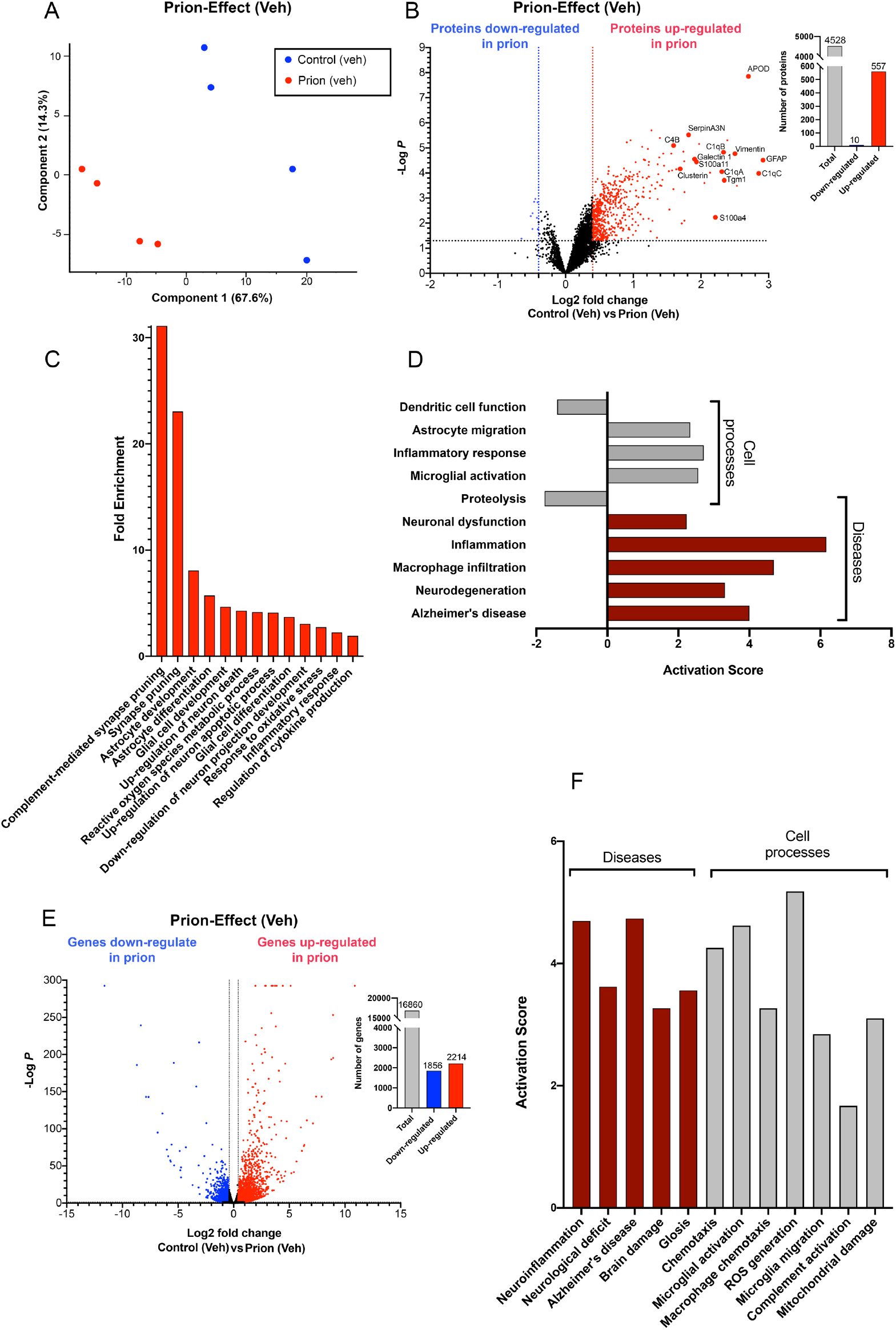
Markers of neurodegeneration and neuroinflammation that map to those in Alzheimer’s disease are up-regulated in murine prion disease. (**A**) Principal components analysis (PCA) of the global proteomic study of 4 control vehicle (Veh) treated mice and 4 prion-infected animals treated with vehicle. (**B**) Volcano plot of differential expression of proteins in the control-vehicle versus prion-vehicle. Red and blue points represent the proteins which significantly increased or decreased expression, respectively, in prion-vehicle compared to control-vehicle mice (FDR<0.05, ±Log_2_ 0.4-fold change). The bar graph summarises the total number of proteins analysed (grey) and the number of proteins that were significantly increased (red) or decreased (blue) in prion-vehicle compared to control-vehicle mice (FDR<0.05, ±Log_2_ 0.4-fold change). (**C**) Gene ontology ((GO) analysis of proteins that are significantly up-regulated in prion-vehicle compared to control-vehicle mice. GO ‘Biological process’ terms are plotted against the fold enrichment relative to the expected number of gene lists of these sizes. (**D**) *Pathway Studio* analysis of *cell processes* (grey) and *diseases* (red) associated with the proteins that are significantly up-regulated in prion-vehicle compared to control-vehicle mice. *Pathway Studio* assesses proteomic changes in the input data with functions reported in the literature to assign a directionality index called “activation score”. This indicates the direction of the effect of the proteins on cell processes and diseases. (**E**) *Prion-effect* volcano plot generated using DESeq2 differential gene expression analysis of genes in the prion-vehicle vs control-vehicle. Red and blue points represent genes that are significantly increased or decreased in transcription, respectively, in prion-vehicle compared to control-vehicle mice (FDR<0.05, ±Log_2_ 0.4-fold change). The bar graph summarises the total number of genes analysed (grey) and the number of genes that were significantly increased (red) or decreased (blue) in prion-vehicle compared to control-vehicle mice (FDR<0.05, ±Log_2_ 0.4-fold change). (**F**) Activation scores from the *Pathway Studio* analysis of *cell processes* (grey) and *diseases* (red) associated with the genes that are significantly up-regulated in prion-vehicle compared to control-vehicle mice.

**Table 1.** List of primary antibodies used in western blotting.

| **Antibody** | **Source** | **Dilution** | **Company** | **Catalogue No.** |
| --- | --- | --- | --- | --- |
| GFAP | Mouse | 1:5000 | Sigma | G3893 |
| Vimentin | Mouse | 1:2000 | R n D Systems | MAB21052 |
| APOE | Rabbit | 1:3000 | Abcam | ab183596 |
| SERPINA3N | Mouse | 1:2000 | R n D Systems | AF4709 |
| Galectin-1 | Rabbit | 1:1000 | Sigma | SAB5500110 |
| Clusterin | Mouse | 1:2000 | R n D Systems | AF2747 |
| Prion | Mouse | 1:2000 | Abcam | ab61409 |
| Alpha tubulin | Mouse | 1:5000 | Abcam | ab7291 |

These data point to the adaptive responses in murine prion disease mirroring many of the hallmarks of AD and other human neurodegenerative diseases associated with the propagation of misfolded protein. This correlation was further evident in bioinformatic analysis using *Pathway Studio* that quantitatively assesses proteomic (and transcriptomic) changes with functions reported in the literature to assign an *“activation score”*. In this analysis cell processes such as astrocyte migration, microglial activation and inflammation are seen to have a positive activation score, indicating that these processes are up-regulated, whilst other processes including dendritic cell function and proteolysis are negatively regulated **(Fig 2D, Supplementary Table S4)**. Importantly, assessment of *“diseases caused”* show that the proteomic changes associated with murine prion disease are positively correlated with disease indicators for AD, neurodegeneration and neuronal dysfunction **(Fig 2D)**.

The overlap between the hallmarks of murine prion disease and human neurodegenerative disease were further evident in the transcriptomic analysis of hippocampi from prion diseased animals. In these experiments groups of three control- or prion infected animals were treated with vehicle from 7-11 w.p.i after which hippocampi were dissected and RNA extracted and sequenced **(Supplementary Fig S2A,B).** Of the 24,062 genes covered in the RNAseq ∼17,000 gene transcripts were identified **(Supplementary Table S5 – for total RNAseq data)** of which 2,214 were significantly (p<0.05) up-regulated by more than 0.4 log2 fold (red dots in **Fig 2E, Supplementary Table S6**) and 1,856 significantly down-regulated (blue dots in **Fig 2E**) in prion diseased animals compared to control infected animals. In line with the proteomic studies these experiments revealed an up-regulation of neuroinflammatory pathways including activation of microglia and astrocytes as well as the activation of the complement system as evidenced by positive activation scores derived from *Pathway Studio* **(Fig 2F, Supplementary Table S7)**. Furthermore, oxidative stress and mitochondrial dysfunction, common features of AD ^34, 35^ were also evident in prion disease where positive activation scores for reactive oxygen species (ROS) associated pathways and the up-regulation of markers of mitochondrial damage were observed **(Fig 2F)**. Together, the transcriptomic and proteomic studies demonstrated that murine-prion disease recapitulates many of the key pathological features of AD, including up-regulation of misfolded protein clearance mechanisms (e.g. ABCA1, ApoE), neuroinflammation (astrocyte, microglial and complement activation) and mitochondrial dysregulation. These data support the notion that prion disease, like other neurodegenerative diseases that progress via *“prion-like”* propagation of misfolded protein, share common adaptive mechanisms ^36, 37^.

### M_1_-PAMs *“normalise”* brain processes that are dysregulated in prion disease

Next, we wanted to assess the impact of VU846 administration on the *prion-effect*. In these experiments infected mice were treated daily with VU846 (10mg/kg, i.p.) from 7-10 w.p.i and hippocampi dissected and analysed to assess the status of the global proteome **(Fig 1G).** In contrast to prion diseased animals treated with vehicle where >500 proteins were up-regulated compared to control mice **(Fig 2B)**, only 248 proteins were significantly (p<0.05) up-regulated by more than 0.4 log2 fold in prion mice chronically administered VU846 **(Fig 3A,B, Supplementary Table S8)**. Of those proteins that were up-regulated the extent of the fold change was significantly (p<0.05) less than seen in the vehicle treated animals (**compare Fig 2B with Fig 3B**). These data point to a dampening of the *prion-effect* in animals treated with VU846, an observation reflected in the principal component analysis which indicated that there was little variation between non-infected vehicle treated animals and prion animals treated with VU846 **(Fig 3A).**

**Figure 3.**
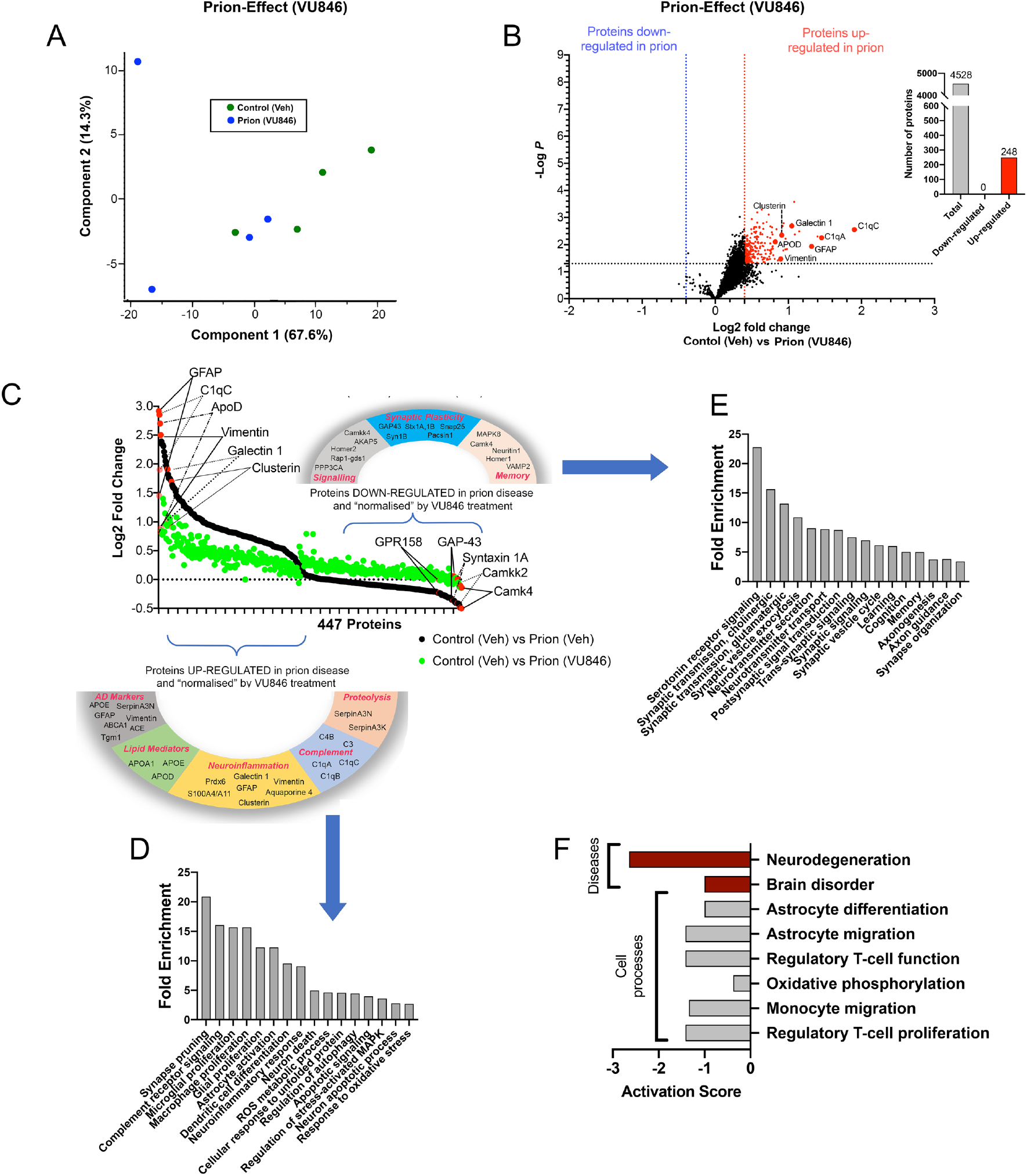
M1-receptor PAM, VU846 *“normalises”* brain processes that are dysregulated in prion disease. (**A**) Principal components analysis (PCA) of the global proteomic study of 4 control vehicle (Veh) treated mice and 4 prion-infected animals treated with VU846. (**B**) Volcano plot representation of differential expression of proteins in the control-vehicle vs prion-VU846. Red and blue points represent the proteins that were significantly increased or decreased in expression, respectively, in prion-VU846 compared to control-vehicle mice (FDR<0.05, ±Log_2_ 0.4-fold change). Bar graph represents the total number of proteins analysed (grey) and the number of proteins that were significantly up-regulated (red) or down-regulated (blue) in control-vehicle vs prion-VU846 (FDR<0.05, ±Log_2_ 0.4-fold change). (**C**) *‘Normalisation plot’* comparing the *prion-effect* vehicle to *prion-effect* VU846. The 477 proteins that were significantly different (*P*<0.05) between these two groups were used to generate the plot with black dots representing *prion-effect* in animals treated with vehicle and green dots representing *prion-effect* in animals treated with VU846. (**D**) GO analysis of proteins that are significantly up-regulated in the *prion-effect* vehicle and ‘normalised’ in the *prion-effect* VU846. The GO ‘biological process’ terms are plotted against the fold enrichment relative to the expected number of gene lists of these sizes. (**E**) GO analysis of proteins that are significantly down-regulated in the *prion-effect* vehicle and ‘normalised’ in the *prion-effect* VU846. The GO ‘biological process’ terms are plotted against the fold enrichment relative to the expected number of gene lists of these sizes. (**F**) *Pathway Studio* analysis of the overall impact of VU846-mediated normalisation of proteins that are either up- or down-regulated in prion disease. The *cell processes* are shown in grey, and *disease processes* in red.

As a control for these experiments, we assessed changes to the hippocampal proteome in animals that were not infected with prion but administered with vehicle or VU846. Remarkably, under these conditions there was very little effect of drug with only 42 significantly up-regulated proteins and no proteins showing down-regulation **(Supplementary Fig S3, Supplementary Table S9)**.

We extended this analysis to examine proteins that showed a significant difference between the *prion-effect* in animals treated with vehicle (black dots in **Fig 3C**) vs the *prion-effect* in animals treated with VU846 (green dots in **Fig 3C, Supplementary Table S10**). This analysis revealed 447 proteins that were significantly different (p<0.05) between these two groups (i.e. *prion-effect* vehicle *vs prion-effect* VU846). Plotting these data generated what we refer to as a “normalisation plot” where the closer the green dots (representing proteins that change in the drug treated animals) were to zero the closer those proteins are to normal, non-disease, levels of expression (Fig 3C).

The normalisation plot illustrates that despite markers of activation of microglia and astrocytes such as GFAP, vimentin, clusterin, and galectin-1, as well as components of the complement system, including C1qA, B, C and the complement receptors C4B and C3, being significantly up-regulated in prion diseased animals treated with VU846, the level of expression of these proteins is less than that seen in prion animals treated with vehicle **(Fig 3C).** Furthermore, markers of AD including the apolipoproteins ApoE, ApoD and ApoC ^33, 38, 39^ as well as key enzyme markers such as Tgm-1 ^40, 41^ and regulators of proteolysis (e.g. serpinA3N and serpinA3K) ^42, 43^, which in the context of prion disease were all up-regulated, showed lower levels of expression in prion animals treated with VU846 **(Fig 3C).** GO enrichment analysis indicated that those proteins showing lower levels of expression in VU846 fell into groups associated with disease responses such as neuroinflammatory response, apoptotic and neuronal death pathways, ROS metabolism and synapse pruning **(Fig 3D, Supplementary Table S11)** consistent with the observation that VU846 reduced prion disease severity.

Whilst the *prion-effect* in vehicle treated animals was characterised by an up-regulation of neurodegeneration disease indicators (black dots, left hand side of **Fig 3C**) there is also a significant down-regulation of proteins associated with synaptic function (black dots, right hand side of **Fig 3C, Supplementary Table S10**). Hence, synaptic proteins including SNAP-25 and syntaxin 1A/1B as well as signalling proteins including calcium-calmodulin protein kinase 4 (Camk4) and mitogen-activated protein kinases (e.g. MAPK8) had decreased expression in prion disease **(Fig 3C).** Remarkably, these proteins were expressed at near normal levels in prion diseased animals treated with VU846 **(Fig 3C, Supplementary Table S10)** suggesting that processes identified in the GO analysis of these proteins such as synaptic organisation, memory, neurotransmission secretion and long-term potentiation are disrupted in prion disease and *“normalised”* by treatment with VU846 **(Fig 3E, Supplementary Table S11)**. In this way key brain functions lost in prion disease might be preserved by treatment with VU846, providing a mechanistic framework for the improved behavioural performance of prion infected animals treated with VU846 (**Fig 1D**).

Finally, the overall impact of VU846-mediated normalisation of proteins that are either up or down-regulated in prion disease was assessed using *Pathway Studio*. The protein expression changes associated with M_1_-PAM treatment generated negative activation scores (i.e. a down-regulation) for inflammatory and mitochondrial cell processes as well as negative scores for diseases such as neurodegeneration **(Fig 3F, Supplementary Table S12),** providing further mechanistic support for the observation that VU846 reduces disease severity.

### Further assessment of the effect of VU846 in prion disease

An alternative approach to the analysis of the effect of VU846 administration on the global proteome in murine prion disease was to directly compare the relative expression levels of proteins in prion diseased animals treated with vehicle vs VU846 **(Fig 4A)**. This comparison we refer to as the *PAM-effect* (i.e. comparing prion-vehicle vs prion-VU846). In this analysis, of the 4,528 proteins quantified, 108 are down-regulated by VU846 treatment and 11 are up-regulated **(Fig 4A, Supplementary Table S13).** Comparing the *PAM-effect* with the *prion-effect* (vehicle) we are able to identify those proteins that are up-regulated in prion disease (*prion-effect*) and determine if they overlap with those proteins that are down-regulated by VU846 (*PAM-effect*) **(Fig 4B, Supplementary Table S14)**. Consistent with the *“normalisation”* analysis above, this approach indicated that many of the up-regulated proteins associated with adaptive processes in prion disease, including those involved in neuroinflammation and the processing of misfolded proteins are subsequently down-regulated in the *PAM-effect*.

**Figure 4.**
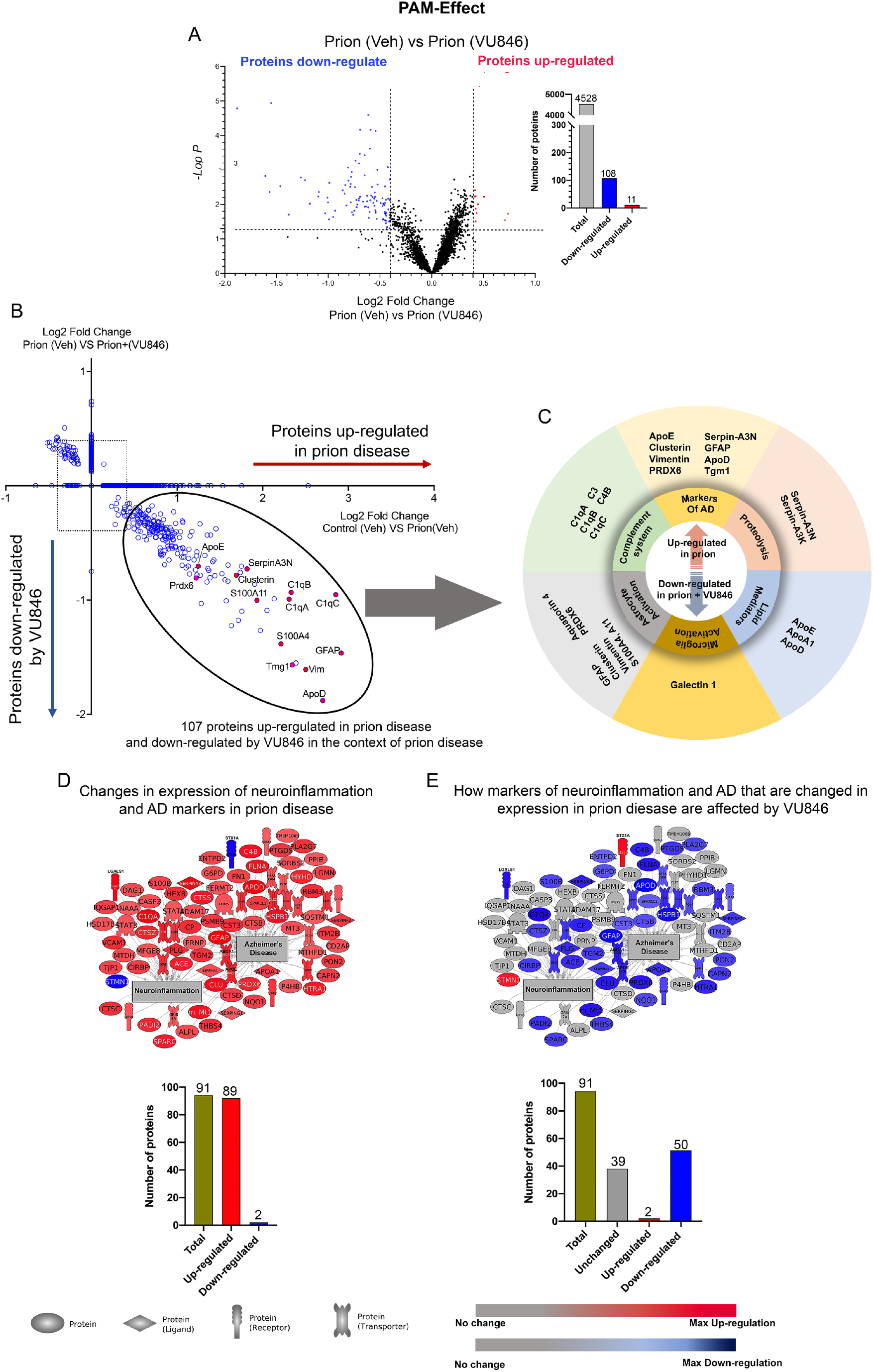
M1-receptor PAM, VU846 has a ‘PAM-effect’ on the prion mouse hippocampal proteome. (**A**) Volcano plot of differential expression of proteins in prion-VU846 vs prion-vehicle (Veh), the so-called *PAM-effect*. Blue and red points represent the proteins that are significantly decreased or increased in expression, respectively, in prion-VU846 compared to prion-vehicle mice (FDR<0.05, ±Log_2_ 0.4-fold change). Bar graph represents the total number of proteins analysed (grey) and the number of proteins that were significantly down-regulated (blue) or up-regulated (red) in prion-VU846 vs prion-vehicle (FDR<0.05, ±Log_2_ 0.4-fold change). (**B**) Quadrant scatter plot showing the effect of VU846 in the context of prion disease. The x-axis shows fold changes of proteins that are up- or down-regulated by prion disease (comparison between control-vehicle and prion-vehicle; *prion-effect*) and the y-axis depicts fold changes of proteins that are up- or down-regulated by VU846 in prion disease (comparison between prion-VU846 and prion-vehicle; *PAM-effect*). The expression of proteins lying on the x-axis are changed by prion disease but not by VU846, whereas the expression of proteins lying on the y-axis are not changed in prion disease but are changed by VU846. Proteins outside the square box are significantly changed in expression (FDR<0.05, ±Log_2_ 0.4-fold change). (**C**) Grouping of proteins associated with neurodegenerative disease that are up-regulated in the *prion-effect* and down regulated in the *PAM-effect*. Representative image generated with *Pathway Studio* showing 94 proteins from the proteomic dataset whose overall expression levels are associated with promoting Alzheimer’s disease and neuroinflammation. The bar graph summarises these changes. Of these 94 proteins (green), 92 are up-regulated by *prion-effect* (prion-vehicle versus control vehicle) (red) and only 2 proteins are down-regulated (blue). Effect of VU846 on proteins that are shown to promote Alzheimer’s disease. The bar graph summarises these changes. Of these 94 proteins (green), 92 proteins were up-regulated by *prion-effect* (red) 54 were down-regulated by the *PAM-effect* (blue) whereas the 2 proteins up-regulated by *prion-effect* are down-regulated by VU846 (red). 38 proteins remain unchanged (grey).

Further analysis using *Pathway Studio* identified those proteins associated with neuroinflammation and AD that are changed in prion disease (i.e. the *prion-effect* in vehicle treated animals **(Fig 4D, Supplementary Table S15).** Taking these same proteins, we asked which are changed in the *PAM-effect* (i.e. prion-vehicle vs prion-VU846). We found that over half of the 94 proteins associated with neuroinflammation/AD in the *prion-effect* are changed in the *PAM-effect* **(Fig 4D,E, Supplementary Table S15).** Importantly, the direction of change seen in the *PAM-effect* is opposite to that seen in the *prion-effect*. Hence, those proteins that are down-regulated in the *PAM-effect* (blue in **Fig 4E**) are up-regulated in the *prion-effect* (red in **Fig 4D**). It is equally important to note that only a proportion (∼60%) of the proteins associated with neuroinflammation/AD in the *prion-effect* are subsequently changed in the *PAM-effect* (**Fig 4E).** Hence, VU846 appears to mediate a partial correction of the dysregulated proteins associated neuroinflammation/AD that likely contributes to the observation that PAM treatment slows but does not completely halt disease progression (**Fig 1E**).

### Validation of the impact of VU846 in prion disease – biomarkers of disease modification

We selected key indicators of VU846 activity emerging from the mass spectrometry studies to probe in western blots. These consisted of markers of astrocyte and microglial activation (GFAP, vimentin, galectin-1 and clusterin) and indicators of neurodegenerative disease (ApoE and serpinA3N) that each demonstrated significant *prion-effects* and *PAM-effects* from the mass spectrometry proteomic analysis **(Fig 5A-F).** Consistent with the mass spectrometry data all the protein markers tested were up-regulated in expression in vehicle treated prion diseased animals and normalised to non-infected animal levels in prion diseased animals treated with VU846 **(Fig 5G and H).** These experiments not only confirmed the mass spectrometry data, they established that western blotting could be used in future studies to probe the disease modifying properties of M_1-_receptor ligands.

**Figure 5.**
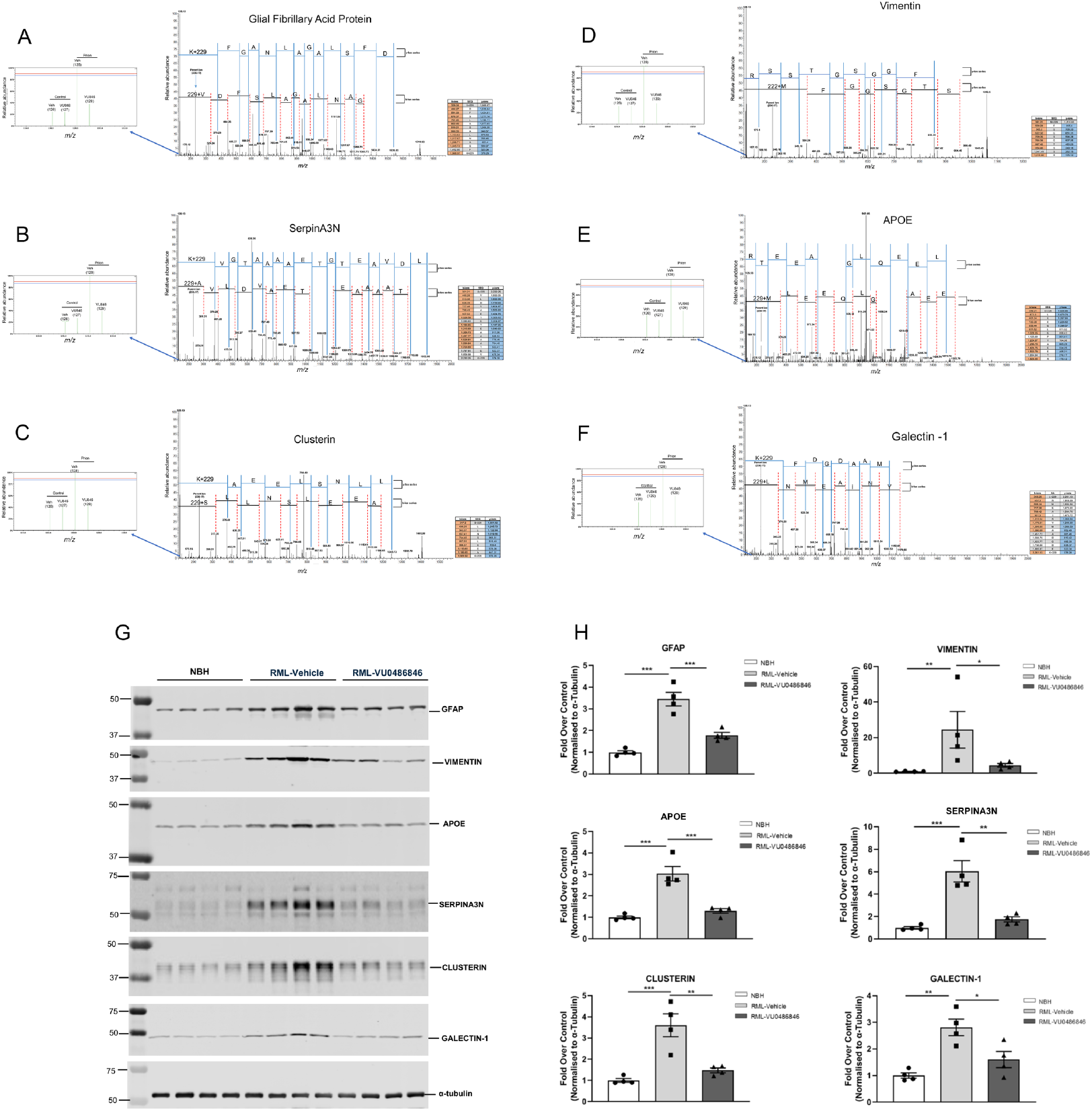
Markers of disease modification modulated by VU846 are validated by western blotting. (**A-F**) Representative spectra from the mass spectrometry-proteomics data shows that markers of astrocyte and microglial activation and neurodegenerative disease are significantly modified by the *prion-effect* and *PAM-effect*. These include (**A**) GFAP, (**B**) SerpinA3N, (**C**) clusterin, (**D**) vimentin, (**E**) ApoE and (**F**) galectin-1. (**G**) Western blot of hippocampi showing changes in protein expression of the selected markers in the control mice treated with vehicle and prion- infected mice treated with vehicle or VU846 (10 mg/kg). Each lane represents a different mouse. The primary antibodies used are detailed in **Table 1**. (**H**) Quantification of western blots shown in G. Data are shown as mean ± SEM, with individual data points displayed within the bars. n=4.

### Different M_1_-PAMs mediate similar effects in prion disease

We next tested if the prototypical M_1_-PAM, BQCA, that we have previously revealed could restore learning and memory deficits in prion disease and slow disease progression ^14^, had similar effects on the proteome of prion infected mice as that observed for VU846. This was a smaller scale experiment **(see; methods and Supplementary FigS4)** where 2,202 proteins qualified for analysis and of these 56 were up-regulated and 15 down-regulated in the *prion-effect* in vehicle treated animals **(Supplementary FigS4, Supplementary Table S16)**. This *prion-effect* was dampened by chronic daily treatment of BQCA from 7 w.p.i **(Supplementary FigS5B, Supplementary Table S17).** Importantly, the proteins regulated by BQCA fell into the same classes of proteins that were regulated by VU846; those involved in neuroinflammation (e.g. GFAP, vimentin, galectin 1, clusterin) and markers of AD including, ApoE, ApoO, S100 proteins and Prdx6 **(Supplementary FigS5B, Supplementary Table S17).** Thus, two chemically distinct M_1_-PAMs (BQCA and VU846) similarly impacted neuroinflammatory and disease-adaption processes.

### Transcriptomic studies reveal additional prion disease modification by an **M_1_**-receptor PAM

To complement the proteomic studies, we conducted a global transcriptomic analysis of hippocampi derived from animals treated daily with VU846 from 7-11 w.p.i. **(Supplementary Fig S2A).** The transcriptomic *prion-effect* in vehicle treated animals consisted of >1,800 genes that were down-regulated and >2,200 genes that were up-regulated **(Fig 2E)**. This *prion-effect* was substantially dampened in animals treated with VU846 where only 168 genes were down-regulated and 888 up-regulated in prion infected animals treated with VU846 compared to drug treatment of non-infected controls **(Fig 6A, Supplementary Table S18).** In contrast, comparing control animals treated with vehicle vs control animals treated with VU846 we observed little effect of VU846 **(Supplementary Fig S6, Supplementary Table S19).**

**Figure 6.**
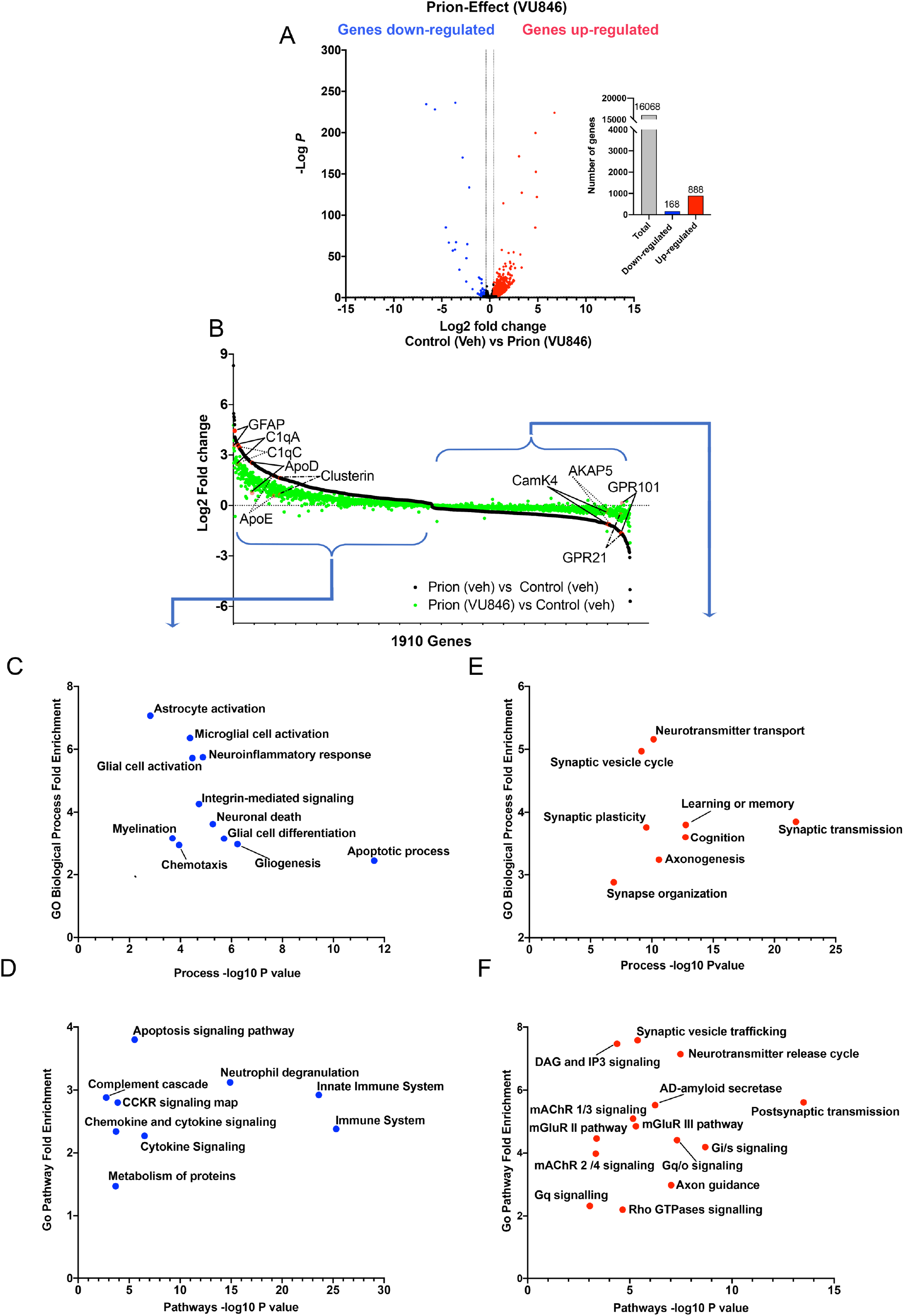
Transcriptomic studies reveal additional prion disease modification by VU846. (**A**) Volcano plot of genes in the control-vehicle (Veh) versus prion-VU846. The plot was generated using DESeq2 differential gene transcription in prion-VU846 treated mice compared to prion-vehicle mice. Red and blue points represent genes with significantly increased or decreased levels of transcript, respectively, in prion-VU846 compared to control-vehicle mice (FDR<0.05, ±Log_2_ 0.4-fold change). Of the 16,068 genes analysed, (grey bar), 888 genes were up-regulated (red bar) and 168 genes were down-regulated (blue bar). (**B**) ‘Normalisation plot’ comparing the *prion-effect* vehicle to *prion-effect* VU846. The 1,910 genes that were significantly different (*P*<0.05) between these two groups were used to generate the plot, with black dots representing *prion-effect* in animals treated with vehicle and green dots representing *prion-effect* in animals treated with VU846. (**C,D**) Fisher exact test for (**C**) GO ‘Biological processes’ and (**D**) GO ‘Pathways’ of the genes that are significantly up-regulated in the *prion-effect* vehicle and ‘normalised’ in the *prion-effect* VU846. The x-axis is the -log_10_ of the *P* value obtained from the Fisher exact test and the y-axis is the relative difference between the percentages of significantly differentially transcribed genes that carried the depicted annotations over the percentage of all sites that carried the same annotation. **(E,F)** The Fisher exact test for (**E**) GO ‘Biological processes’ and (**F**) GO ‘Pathways’ of the genes that are significantly down-regulated in the *prion-effect* vehicle and ‘normalised’ in the *prion-effect* VU846.

In analysis similar to that conducted in the proteomic study, we constructed a *“normalisation plot”* of the genes that were differentially transcribed between the *prion-effect* observed in vehicle treated animals (black dots in **Fig 6B, Supplementary Table S20**) and the *prion-effect* observed in VU846 treated animals (green dots in **Fig 6B, Supplementary Table S20**). GO Biological Process and Pathway analyses established that genes that were up-regulated in prion disease but that were significantly less up-regulated in VU846 treated animals (left hand-side of **Fig 6B, Supplementary Table S21**) were associated with neurodegenerative disease processes. These included indicators of neuroinflammation such as astrocyte and microglial activation as well as genes associated with neuronal death and apoptosis **(Fig 6C, Supplementary Table S21)**. Furthermore, consistent with the notion that neuroinflammatory pathways were activated in prion disease and diminished by VU846, the GO Pathway analysis revealed enrichment of genes in pathways such as the complement cascade and chemokine/cytokine signalling **(Fig 6D, Supplementary Table S21)**. Conversely, a group of genes that were down-regulated in prion disease were transcribed at significantly higher levels in the *prion-effect* of animals treated with VU846 (right hand-side of **Fig 6B**). These genes were enriched in brain processes that are known to be disrupted in neurodegenerative disease, including synaptic plasticity, learning and memory, cognition and synaptic transmission **(Fig 6E, Supplementary Table S21)**. GO Pathway analysis also revealed enrichment in pathways associated with neuronal activity including muscarinic and glutamate receptor signalling, synaptic vesicle trafficking and G protein signalling **(Fig 6F, Supplementary Table S21)**. Overall, these transcriptional data are consistent with the proteomic studies in establishing that treatment with VU846 corrects, or normalises, dysregulated gene transcription associated with prion disease; reducing the transcription of genes associated with disease whilst increasing gene transcription associated with normal brain function and synaptic activity.

We next assessed the changes in gene transcription that are associated with the *PAM-effect* **(Fig 7A, Supplementary Table S22)**. Here we observed that VU846 treatment resulted in the transcriptional down-regulation 877 genes and the up-regulation of 712. Analysis of these changes in *Pathway Studio* revealed negative activation scores for neuroinflammation, complement activation and chemotaxis consistent with an anti-inflammatory response to VU846 treatment **(Fig 7B, Supplementary Table S23)**. Furthermore, negative activation scores for neurological diseases including AD and mild cognitive impairment were observed in the global analysis of the *PAM-effect* **(Fig 7B)**. These data pointed to a reduction in disease severity in VU846 treated animals, an observation further supported by plotting the *prion-effect* in vehicle treated mice with the *PAM-effect*. This revealed 817 genes that were transcriptionally up-regulated in the *prion-effect* and subsequently down-regulated in the *PAM-effect* (lower right-hand quadrant in **Fig 7C, Supplementary Table S24**). Among these are genes associated with the complement system, microglia and astrocyte activation and included genes for proteins that were similarly regulated in the proteomic analysis described above (green text in **Fig 7D**). There was a similar overlap between the proteomic and transcriptomic data sets in markers of AD, proteolysis and lipid mediators **(Fig 7D)**. Whereas not all transcriptional changes were linked with corresponding proteomic changes, the combined data sets strongly supported the notion that VU846 reduces neuroinflammation and markers of neurodegenerative disease.

**Figure 7.**
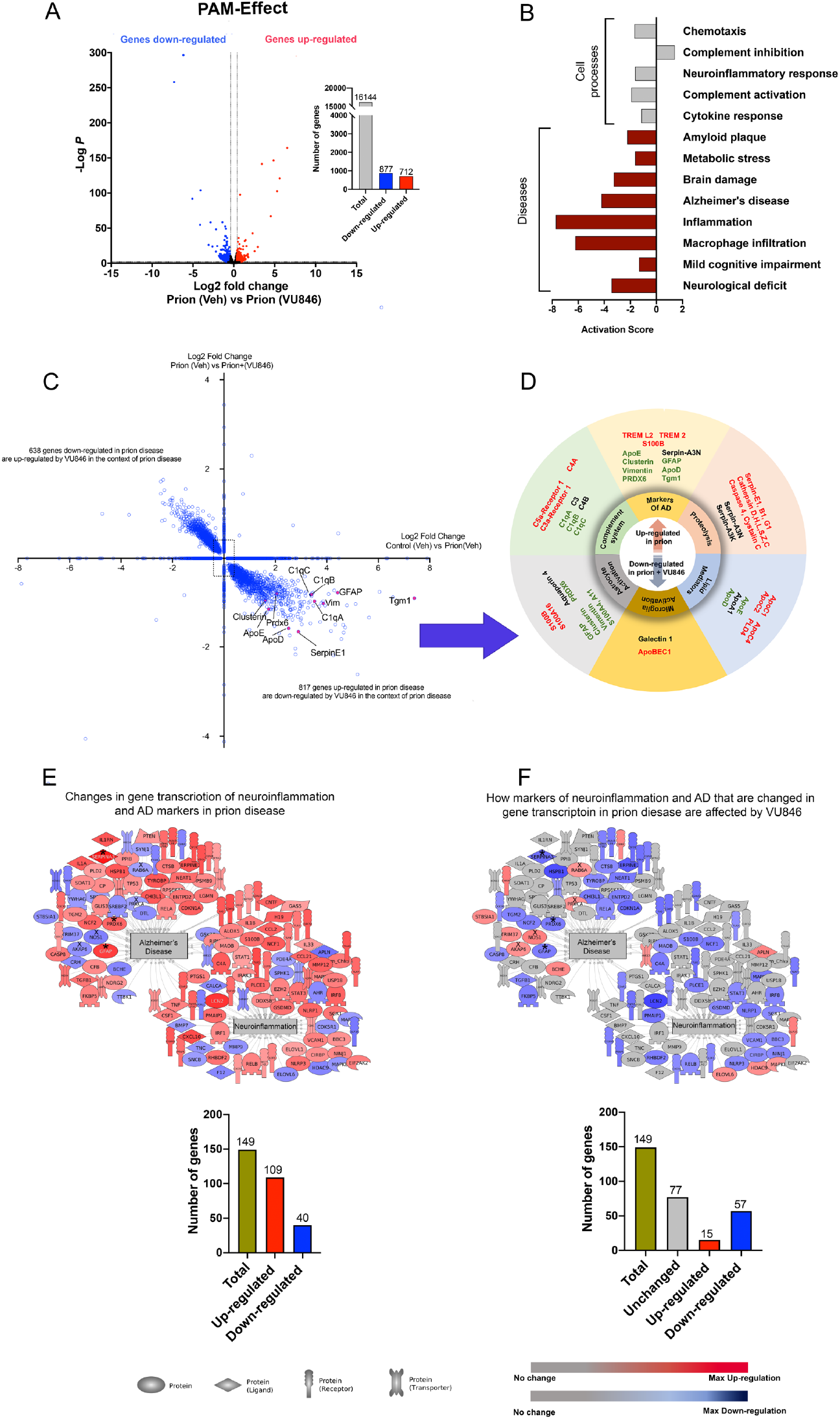
M1-receptor PAM, VU846 has a ‘*PAM-effect’* on the prion mouse hippocampal transcriptome. (**A**) Volcano plot from DESeq2 differential gene transcription of VU846 in the context of prion disease. The plot was generated using DESeq2 differential gene transcription prion-VU846 treated mice compared to prion-vehicle mice. Of the 16,144 genes analysed (grey bar) 712 genes were up-regulated (red bar) and 877 genes were down-regulated (blue bar) by prion-VU846 compared to prion-vehicle. Red and blue points represent genes with significantly increased or decreased transcription respectively in prion-VU846 compared to prion-vehicle mice (FDR<0.05, ±Log_2_ 0.4-fold change). (**B**) *Pathway Studio* analysis of *cell processes* (grey) and *diseases* (red) associated with the genes that are significantly up- and down-regulated in prion-VU846 compared to prion-vehicle. (**C**) Quadrant scatter plot showing the effect of VU846 on gene transcription in the context of prion. The x-axis shows fold changes of genes that are up- or down-regulated by prion disease (comparison between control-vehicle and prion-vehicle; *prion-effect*) and the y-axis depicts fold changes of proteins that are up- or down-regulated by VU846 in prion disease (comparison between prion-vehicle and prion-VU846; *PAM-effect*). (**D**) Grouping of genes associated with neurodegenerative disease that are up-regulated in the *prion-effect* and down regulated in the *PAM-effect*. Genes that are identified in the proteomic analysis are indicated in black text, those identified in the transcriptomic analysis indicated in red text and those identified in both proteomic and transcriptomic data indicated in green text. (**E-F**) Representative images generated with *Pathway Studio* showing the link between VU846 treatment and regulation of neuroinflammation and markers of AD. A proportion of the genes that are up- or down-regulated in the *prion-effect* are subsequently affected by VU846 in the *PAM-effect*.

The transcriptomic link between VU846 treatment and regulation of neuroinflammation and markers of AD was further probed using *Pathway Studio.* In this analysis we identified all the gene changes associated with neuroinflammation and AD in the *prion-effect* in vehicle treated animals **(Fig 7E, Supplementary Table S25).** We then asked, “which of these genes are also affected by drug in the *PAM-effect* (marked in blue/red in **Fig 7F, Supplementary Table S25**)?” The outcome of this analysis was that only a proportion (∼48%) of the genes that are up or down-regulated in prion disease are subsequently affected by VU846 in the *PAM-effect*. Importantly, of those genes that are changed in the *PAM-effect*, the direction of the effect is opposite to that seen in the *prion-effect*. For example, GFAP, Prdx6 and serpinA3 are all up-regulated in the *prion-effect* (marked with an asterix in **Fig 7E**) and these same genes are down-regulated in the *PAM-effect* (**Fig 7F**). In contrast, genes encoding signalling proteins such as protein kinase C*α* (PRKCA), RAB6A, NOS1 and AKAP6 are down-regulated in the prion-effect (marked with a cross in **Fig 7E**) but subsequently up-regulated in the PAM-effect (**Fig 7F**).

## DISCUSSION

All attempts to identify a therapy that can substantially delay the progression of neurodegenerative disease, including AD, have so-far failed in preclinical development or clinical trials ^6, 44, 45^. The emergence of an array of AD risk factors from GWAS ^2–4^ has provided a number of potential novel targets that are distinct from the extensively tested *β*- and *γ*-secretase inhibitors and antibodies that target abnormal A*β* ^6, 44, 45^. However, the paucity of knowledge of how these proteins operate in the context of neurodegenerative disease, and the intractability of many of these as pharmacological targets, has limited pre-clinical validation and subsequent drug-discovery efforts ^46, 47^. Here we describe how targeting the M_1_-receptor with PAMs that amplify the spatial and temporal patterns of physiological stimulation of M_1_-receptors by acetylcholine can reduce the emergence of neuroinflammation and adaptive processes associated with prion-mediated neurodegeneration. Furthermore, our proteomic and transcriptomics data point to such M_1_-receptor activation as critical for maintaining synaptic function and mitochondrial/redox homeostasis. In this way our study provides evidence for the M_1_-receptor as an attractive therapeutic target that both can both reduce neurodegenerative disease severity and maintain synaptic function, thereby increasing life-span and maintaining normal behaviour.

The muscarinic receptor family members were among the first GPCRs to be cloned and characterised and, as such, considered by many as prototypical ^48^. The ensuing decades of research have resulted in a rich understanding of the signalling, pharmacology and physiological roles of this receptor family ^49, 50^. The M_1_-receptor in particular with its high expression in memory centres and pro-cognitive properties has been proposed as a target for the treatment of memory loss in AD that would avoid the dose-limiting adverse responses associated with current clinically approved cholinesterase inhibitors ^23, 24^. The challenge has been to develop M_1_-receptor selective drugs since the orthosteric acetylcholine binding site is highly conserved across the five muscarinic receptor subtypes ^20^. As such orthosteric ligands, such as xanomeline, have failed in the clinic as AD therapeutics primarily due to cholinergic-adverse responses mediated by peripheral M_2_- and M_3_-receptors ^21^. An alternative strategy has been to target the allosteric pocket through ligands that act in a receptor-subtype selective manner to enhance receptor activity in a co-operative manner with the natural ligand acetylcholine ^24^. The prospect that these PAMs might offer an approach to the treatment of memory loss in AD by the restoration of cholinergic transmission has led to an expansion of PAM-chemotypes displaying various pharmacological profiles ^18, 51, 52^. This has allowed our laboratories to interrogate the preclinical pharmacology of PAMs with distinct levels of intrinsic efficacy, biased agonism, and levels of cooperativity with the physiological agonist, acetylcholine. These studies led to the conclusion that M_1_-receptor PAMs that display moderate to high levels of co-operativity with acetylcholine whilst having low intrinsic efficacy and no ligand-bias would provide pro-cognitive effects with little associated cholinergic adverse responses ^18, 53^. VU846 shows many of these favoured characteristics and together with good brain penetration and favourable DMPK properties ^27^ makes for an excellent proof-of-concept M_1_-receptor selective PAM to broadly assess the pre-clinical benefits of this class of pharmacological agent.

Murine prion disease is associated with a disruption in hippocampal cholinergic innervation that results in defective learning and memory that can be restored with clinically approved cholinersterase inhibitors and the orthosteric muscarinic agonist xanomeline ^14^. We reasoned therefore that murine prion disease was a good model to investigate novel therapeutic approaches to restore defective cholinergic transmission and subsequently demonstrated that the M_1_-PAM, BQCA, could similarly restore defective learning and memory in this mouse model ^14^. What was not clear from these previous studies was whether prion disease also exhibited other hallmarks of AD that would allow for a more extended application of this model in preclinical assessment of treatments that could modify neurodegenerative disease progression. By employing global proteomic and transcriptomic analyses we establish here that murine prion disease is associated with neuroinflammation, markers of mitochondrial dysfunction and increased oxidative stress, all of which are also associated with AD and other forms of human neurodegenerative disease ^54–58^. Moreover, we revealed that murine prion disease is associated with the up-regulation of markers of AD and in particular proteins that are involved in misfolded protein clearance, indicating that the disease-adaptive changes potentially associated with compensatory mechanisms in human neurodegenerative are also in operation in murine prion disease. Our study therefore supports the notion that neurodegenerative diseases that are the result of *“prion-like”* spreading and propagation of misfolded protein share many common adaptive and pathological features ^36, 37^.

Given the common features between murine prion disease and human neurodegenerative disease it is particularly noteworthy that chronic treatment with the M_1_-PAM, VU846, resulted in a significant, broad, reduction in neuroinflammation and markers of AD in a manner that correlated with a slowing of disease progression and maintenance of normal mouse behaviour. Importantly, dosing of VU846 commenced at a disease-stage where pathological markers of disease (e.g. accumulation of misfolded PrPsc) were already evident, indicating that VU846 was able to modify disease even when disease had been established. Nonetheless, although many of the markers of neuroinflammation and AD were maintained at lower or near normal levels by VU846 treatment this was not true for all markers. Hence, among the neuroinflammatory and AD markers seen to change in prion disease ∼60% were affected by VU846 treatment. These data point to the possibility that specific features of neuroinflammation and disease-adaptation may be directly regulated by M_1_-receptor activity.

Finally, a striking feature of VU846 activity was that there was little effect of the compound in normal, non-diseased, animals. Hence, it is in a disease context where VU846 has the most profound impact on the proteome and transcriptome. This correlates well with previous reports where M_1_-PAMs have been reported to have little behavioural effect in control animals and it is only when there is a disruption in cholinergic transmission mediated by pharmacological intervention or by neurological disease that M_1_-PAMs are seen to have a significant impact ^14, 27, 59^.

In conclusion, our study reports that treatment of prion-diseased mice with a highly tolerated M_1_-PAM that displays low intrinsic activity and high levels of co-operatively results in a reduction in neuroinflammation and markers of *“AD-like”* disease-adaptive responses. This provides mechanistic insight into the observed slowing of disease progression and maintenance of normal behaviour mediated by the administration of the M_1_-PAM (VU846) and supports the notion that M_1_-PAMs might not only provide a novel therapeutic strategy for the symptomatic treatment of memory loss in AD but also might deliver neuroprotection that extends life span and preserves normal behaviour.

## METHODS

### Animal maintenance

The mice were fed ad libitum with a standard mouse chow. The MloxP Tg37 transgenic mice that overexpresses mouse cellular prion protein (PrP^C^) as described in previous studies ^14, 60^ were provided by Professor Giovanna Mallucci (University of Cambridge Dementia Research Institute). All animal work and care were carried out under a project license according to United Kingdom Home Office Regulations.

### Inoculation of Tg37 mice with prion

Tg37 hemizygous mice at 3 to 4 weeks were inoculated by intracerebral injection into the right parietal lobe with approximately 20 μl of 1% (w/v) brain homogenate of Rocky Mountain Laboratory (RML) scrapie prion as described previously (Mallucci *et al*., 2002; Bradley *et al*., 2017). Control Tg37 mice were inoculated in similar manner with approximately 20 μl of 1% normal brain homogenate (NBH).

### Survival studies

Tg37 mice were inoculated with RML or NBH as described above. The mice were given intra-peritoneal (i.p.) injection of vehicle (20% tween-80), BQCA (15 mg/kg) or VU0486846 (VU846, 10 mg/kg) daily from 7 weeks post-inoculation (w.p.i.). Video recordings of the mice were taken every three days from 7 w.p.i. The mice were examined daily for early indicators and confirmatory signs of scrapie prion disease, and animals were culled when they developed clinical signs of scrapie. Early indicators include clasping of hind legs when mice are lifted by the tail, un-sustained hunched posture, rigid tail, mild loss of coordination, piloerection, and being subdued. Confirmatory signs include sustained hunched posture, ataxia, dragging of limbs, significantly abnormal breathing and impaired writhing reflex. The presence of two early indicator signs plus one confirmatory sign, or two confirmatory signs alone was indication of clinical disease.

### Fear conditioning learning and memory test

The fear conditioning experiments were conducted on male mice at 9 w.p.i. with RML or NBH, prior to the appearance of clinical symptoms. Mice were acclimatised to the behavioural room overnight prior to day of the test. M1 PAM VU846 (10mg/kg) or vehicle were administered via i.p. injection on the day of the behavioural test, 30 minutes prior the training. For fear conditioning, mice were placed in the conditioning chamber (Stoelting ANY-maze Fear Conditioning System, Dublin) and allowed to adapt to the chamber for 2 minutes. The mice received 3 tone/foot shock pairings, where the foot shock (unconditioned stimulus (US); 2 seconds; 0.4 mA) always co-terminated with a tone (conditioned stimulus (CS); 2.8 kH; 85 dB; 30 seconds). The CS-US pairings were separated by 1-minute intervals. After completion of the training, the mice remained in the conditioning chamber for 1 minute and were then returned to their home cages. The next day, the mice were placed back in the conditioning chamber, and time spent immobile was recorded for 3 minutes to assess context-dependent learning. The data were analysed using ANY-maze software (Stoelting, Dublin).

### Burrowing

Assessment of burrowing activity was conducted on mice from 7 w.p.i. to 9 w.p.i. A day prior to the burrowing test, mice were placed into individual burrowing cages containing an empty burrowing tube for a 2-hour period to acclimatise. The burrowing tube is a clear, acrylic tubing with one end sealed with transparent plastic. On the test day, mice received vehicle (20% tween-80) or VU846 (10 mg/kg) via i.p. injection 30 minutes prior to the burrowing test. The mice were placed into individual burrowing cages containing a burrowing tube filled with 140 g of food pellets for 2 hours. The amount of food pellets remaining after the 2 hours was weighed and the burrowing activity was calculated by subtracting the weight of food pellets remaining from the starting weight and expressing the proportion of food pellets that had been displaced as a percentage. The mice were returned to their home cages, and the experiment was repeated on a weekly basis.

### Cortical neuronal primary cultures

Cortical neurons were isolated from 16-day old embryos of C57BL/6 mice. Dissected brains were immediately placed in ice cold dissection buffer (DMEM) and the cerebral cortices were isolated under a dissecting microscope. Cortex tissues were then mechanically triturated, and cells were resuspended in HBSS followed by centrifugation at 500 x g for 5 minutes. The pellets were resuspended in warm neurobasal media supplemented with B-27, L-glutamine and 1% penicillin/streptomycin. The primary cells were plated at a density of 60,000 cells/well in a 96-well microplate that had been pre-coated with 50µg/ml of poly-D-lysine and maintained at 37 ^O^C in a 5% CO_2_ humidified atmosphere. In-vitro assays were performed a week later.

### IP1 accumulation assay

Cells were washed and incubated in 80 μl of 1X stimulation buffer (10 mM HEPES, 1 mM CaCl_2_, 0.5 mM MgCl_2_, 4.2 mM KCl, 146 mM NaCl, 5.5 mM glucose, 50 mM LiCl, pH 7.4) for 1 hour at 37 ^O^C prior drug to treatments. 10 μl of 10X concentrated M1-PAM (BQCA or VU846) was added to respective wells in the microplate, followed by 10 μl of 10X concentrated ACh and incubated at 37 ^O^C for one hour. The stimulation buffer was removed, and cell lysis buffer (IP-One assay kit, CisBio) was added (40 µl/well) and incubated for 10 minutes with shaking at 600 rpm. The cells suspensions (7µl/well) were added to 384-well white ProxiPlates and centrifuged briefly. IP1-d2 conjugate and the anti-IP1 cryptate Tb conjugate (IP-One Tb™ assay kit, CisBio) were diluted 1:40 in lysis buffer and 3 μl of each was added to each well. The plate was incubated at 37 ^O^C for 1 hour and fluorescence resonance energy transfer (FRET) between d2-conjugated IP1 (emission at 665 nm) and Lumi4-Tb cryptate conjugated anti-IP1 antibody (emission at 620 nm) was detected using an Envision plate reader (PerkinElmer). Results were calculated from the 665/620 nm ratio and normalised to the maximum response stimulated by acetylcholine.

### Hippocampal lysate preparation and western blot analysis

Frozen mouse hippocampi were transferred into microcentrifuge tubes containing 300 μl of RIPA buffer [50 mM Tris-HCl, 1 mM EDTA, 1 mM EGTA, 1% (v/v) triton X-100, 0.1% (v/v) 2-mercaptoethanol, pH 7.5] and sonicated three times for 15 seconds each at 3-5 μm amplitude. The lysate was incubated at 4 ^O^C for 2 hours with end-to-end rotation and then centrifuged at 15000 x g for 10 minutes at 4 ^O^C. The supernatant was transferred into new tubes and protein concentration determined using the BCA protein assay, kit according to manufacturer’s instruction (ThermoFisher). 10 μg of protein was added to equal volume of 2X Laemmli sample buffer and heated at 95 ^O^C for 5 minutes and then separated by electrophoresis on a 12% SDS-tris-glycine polyacrylamide gel. The proteins were transferred onto a nitrocellulose membrane, blocked in 5% (w/v) fat-free milk and then immunoblotted with respective primary antibodies overnight at 4 ^O^C. The primary antibodies used in this study are detailed in **Table 1**. After washes and incubation with LI-COR IRDye secondary antibody (LI-COR, Cambridge-UK), the proteins were visualised and quantified using the Empiria Studio software (Li-COR). The intensity of the proteins was normalised to the intensity of alpha tubulin.

### Proteinase K digestion

For proteinase K (PK) digestion analysis, equal volumes of 20 μg/ml of proteinase K and 40 μg of protein lysate were mixed and incubated at 37 ^O^C for 10 minutes. The digestion reaction was stopped by adding Laemmli sample buffer and heating at 95 ^O^C for 5 minutes. The proteins were separated by electrophoresis on a 12% SDS-tris-glycine polyacrylamide gel, and subsequently transferred onto a nitrocellulose membrane and then blocked in 5% (w/v) fat-free milk. The membrane was immunoblotted with anti-prion primary antibody (Abcam, ab61409) overnight at 4 ^O^C, followed by washes and incubation with LI-COR IRDye secondary antibody (LI-COR, Cambridge-UK), the proteins were visualised and quantified using the Empiria Studio software (Li-COR). The intensity of the proteins was normalised to the intensity of alpha tubulin.

### Prion cohorts for proteomics and transcriptomics analysis

For the BQCA proteomics cohort, the mice (NBH and RML) were treated (i.p.) with vehicle (5% glucose) or BQCA (15 mg/kg) daily from 7 w.p.i. for two weeks. Animals were culled and hippocampus dissected. For the VU846 proteomics and transcriptomics cohort, the mice (NBH and RML) were treated (i.p.) with vehicle (20% tween-80) or (10 mg/kg) daily from 7 w.p.i. to 11 w.p.i. Animals were culled, and hippocampus dissected. Hippocampus from one brain hemisphere was processed for mass spectrometry-based proteomics, and the other half for transcriptomics analysis.

### Hippocampal preparation for TMT LC-MS/MS

Mice were killed by cervical displacement and the brain was removed from the skull and dissected immediately. The hippocampi and cortices were flash-frozen on dry ice. The frozen hippocampi (from one hemisphere of the brain) were transferred into microcentrifuge tubes containing SDS lysis buffer (50 mM TEAB, 10% SDS, pH 7.55) supplemented with protease and phosphatase inhibitors and homogenised using a motorised pellet pestle for 60 seconds. 20% CHAPS (w/v) and 10% NP-40 (v/v) were added to final concentrations of 2% and 1% respectively and the lysate was sonicated three times for 15 seconds each at 3-5 μm amplitude and then centrifuged at 15000 x g for 10 minutes at 4 ^O^C. The supernatant was transferred into new microcentrifuge tubes and the protein concentration was determined using the BCA protein assay kit, according to manufacturer’s instruction (ThermoFisher). The lysates were normalised to the same protein concentrations (0.8 mg) with SDS lysis buffer to a final volume 500 μL. The proteins were reduced using 20 mM DTT at 37 ^O^C for 1 hour followed by alkylation in the dark for 30 minutes with 100 mM iodoacetamide. The samples were acidified with 12% phosphoric acid to a final concentration of 1.2% v/v and then digested overnight at 37 ^O^C with sequence grade trypsin at a trypsin-to-protein ratio of 1:20 (w/w) using the ProtiFi S-trap midi digestion columns (ProtiFi, Huntington). Eluted peptides were dried in a vacuum concentrator, resuspended in 0.1% trifluoroacetic acid and desalted using Pierce peptide desalting columns (ThermoFisher). The eluted peptides were dried, resuspended in 50 mM HEPES buffer (pH 8.5) and labelled with TMTSixplex (ThermoFisher) at 25 ^O^C for 2 hours with orbital shaking at 500 rpm. The TMT to peptide ratio was 2.5:1. The labelling reaction was quenched with 5% (v/v) hydroxylamine at final concentration of 0.4% at 25 ^O^C for 30 minutes with orbital shaking at 500 rpm. The peptides were dried, resuspended in 0.1% TFA and separated into ten fractions using the Pierce high-pH reverse-phase fractionation columns (ThermoFisher). The eluted fractions were dried and resuspended in 0.1% formic acid for LC-MS/MS analysis.

For the VU846 cohort, hippocampi from mice in each of the four experimental groups (control + vehicle, control + VU846, prion + vehicle and prion + VU846) were processed, labelled with respective TMT, combined and analysed by LC-MS/MS to give one experimental run. This process was repeated with the other three mice in each experimental group to give four independent runs and datasets (**Figure 1G).** For the BQCA cohort, hippocampi from three mice in each of the experimental groups were pooled together, processed and labelled with respective TMT, combined and analysed by LC-MS/MS to give one experimental run (**Supplementary figure S4).**

### TMT LC-MS/MS and data processing

Samples were analysed by using an LTQ-Orbitrap-Velos mass spectrometer (Thermo Scientific), equipped with an ultra-high-pressure liquid-chromatography system (RSLCnano). The samples were loaded at high-flow rate onto a reverse-phase trap column (0.3 mm i.d. x 1 mm), containing 5 mm C18 300Å Acclaim PepMap medium (Dionex) maintained at 37 ^O^C. The loading buffer was 0.1% formic acid/0.05% TFA/2% ACN. The peptides were eluted from the reverse-phase trap column at a flow rate of 0.3 μl min^-1^ and passed through a reverse-phase PicoFrit capillary column (75 μm i.d. x 400 mm) containing Symmetry C18 100Å medium (Waters) that was packed in-house using a high-pressure device (Proxeon Biosystems). Peptides were eluted over a period of 4 hours, with the output of the column sprayed directly into the nanospray ion source of the LTQ-Orbitrap-Velos mass spectrometer. The LTQ-Orbitrap-Velos mass spectrometer was set to acquire a 1 microscan Fourier transform mass spectrometer (FTMS) scan event at 60,000 resolution over the m/z range of 300–2,000 Da in positive ion mode. The maximum injection time for MS was 500 ms and the AGC target setting was 1e^6^. Accurate calibration of the FTMS scan was achieved using a background ion lock mass for C_6_H_10_O_14_S_3_ (401.922718 Da). Subsequently, up to ten data-dependent higher-energy collision dissociation (HCD) MS/MS were triggered from the FTMS scan. The isolation width was 2.0 Da, with normalized collision energy of 42.5. Dynamic exclusion was enabled. The maximum injection time for MS/MS was 250 ms and the AGC target setting was 5e^4^.

The raw data file obtained from each LC-MS/MS acquisition was processed using Proteome Discoverer (version 2.5.0.400, Thermo Scientific), searching each file in turn using Mascot (version 2.7.07, Matrix Science Ltd.) against the UniProtKB-Swissprot database. The peptide tolerance was set to 10 p.p.m. and the MS/MS tolerance was set to 0.02 Da. A decoy database search was performed. The output from Proteome Discoverer was further processed using Scaffold Q+S (version 4.11.0, Proteome Software). Upon import, the data were searched using X!Tandem (The Global Proteome Machine Organization). PeptideProphet and ProteinProphet (Institute for Systems Biology) probability thresholds of 95% were calculated from the decoy searches and Scaffold was used to calculate an improved 95% peptide and protein probability threshold based on the data from the two different search algorithms.

### Analysis of proteomics data

The data was uploaded in Microsoft Excel (version 2016), Perseus (version 1.6.12.0) and Scaffold (version 4.11.0) analytical suites for downstream analysis. For ease of data handling, all data entries were transformed into log_2_ scale and normalised. For a protein to be included in the analysis, the peptides corresponding to the protein must be present in at least three of the four independent datasets. Contaminants, reverse hits and proteins ‘only identified by site’ were excluded from the analysis. Statistical analyses were performed using 2-tailed Student’s t test, 1-way ANOVA, or 2-way ANOVA. Significance was defined as *P* < 0.05. All statistical tests were performed using GraphPad Prism software. Graphs were plotted using Perseus, Microsoft Excel and GraphPad Prism software.

### Hippocampal preparation for transcriptomics

#### RNA sample preparation

Three mice from each of four experimental groups (control + vehicle, control + VU846, prion + vehicle and prion + VU846) were processed for transcriptomics analysis. RNA from each group was extracted using the RNeasy Plus Mini Kit (Qiagen, Manchester), following the manufacturer’s instructions. The tissue was homogenised in the RNA kit buffer by sonicating three times for 15 seconds each at 3-5 μm amplitude and then centrifuged at 10,000 x g for 3 minutes at 4 ^O^C. The homogenised sample was transferred into the purification columns for RNA purification. RNA was eluted with ultrapure water and concentration and purity measured with NanoDrop-1000 Spectrophotometer (Fisher Scientific).

#### mRNA library construction and data analysis

RNA samples were processed at the Glasgow Polyomics Research Facility. Each sample was subjected to mRNA poly-A enrichment before libraries were generated with TruSeq Stranded mRNA sample preparation kit (Illumina). The libraries were sequenced paired ended (2×75bp) on the NextSeq500 instrument (Illumina) to an average of at least 33 million reads. Raw counts were then converted into FastQC format.

The Galaxy bioinformatics data analysis platform (Version 0.72) was used to process raw data-FastQC files. The data was analysed to remove both the TruSeq3 adaptors used for sequencing and the bad quality RNA sequences using the Trimmomatic flexible read trimming tool for Illumina NGS data (Galaxy version 0.36.5). In particular, transcripts showing 8 hit-read matching any adaptor were trimmed off and discarded, and the sliding window trimming function was applied to eliminate bad quality RNA sequence with a cut-off of 25 phred. The remaining sequences were mapped to the “mouse-mm10” genome using HISAT2, a fast and sensitive alignment program (Galaxy Version 2.1.0) and processed with the StringTie function to assemble and quantify the sequences associated for each gene (BAM files).

Differential gene expression comparisons were performed with “BAM” files using DESeq2 statistical tool (parametric fit type) on the Galaxy bioinformatic platform. Gene differential expression data were analysed with two online software suites; Gene Ontology Panther (http://www.pantherdb.org) and Pathway Studio (www.pathwaystudio.com) to identify diseases, cell processes and pathways associated with genes affected by prion and drug.

### Raw data accession codes

All of the TMT mass spectrometry data, RAW files together with the MaxQuant outputs have been uploaded to PRIDE (Accession code ………).

The raw transcriptomics data have been deposited in the Gene Expression Omnibus repository (Accession code …….).

## Acknowledgements

This work is funded by an MRC Industrial CASE studentship (MR/P016693/1; MS), a University of Glasgow Lord Kelvin Adam Smith Fellowship (SJB), an MRC MICA (MR/P019366/1; ABT, SJB) and a Wellcome Trust Collaborative Award (201529/Z/16/Z; ABT, AC, PMS). We acknowledge Dr Andrew Bottrill the BSU facilities at the Cancer Research UK Beatson Institute (C596/A17196) and the Biological Services at the University of Glasgow.

## Data availability statement

All data are available from the corresponding authors or through the University of Glasgow’s online data repository.

## Author contributions

SJB, ABT Conceived and designed the study and wrote the paper with the aid of the other authors. LD, conducted the proteomic study. MR conducted the transcriptomics study. ARB, conducted mass spectrometry experiments. PH, conducted transcriptomic data analysis. SJB, MS, CM analysis of prion infectivity and conducted animal experiments. SNM provided BQCA. EK, conducted neuronal signalling experiments. PS, AC, designed pharmacology experiments. PJC, CL synthesised experimental compounds and supported data analysis.

## SUPPLEMENTARY FIGURES

**Supplementary figure S1.**
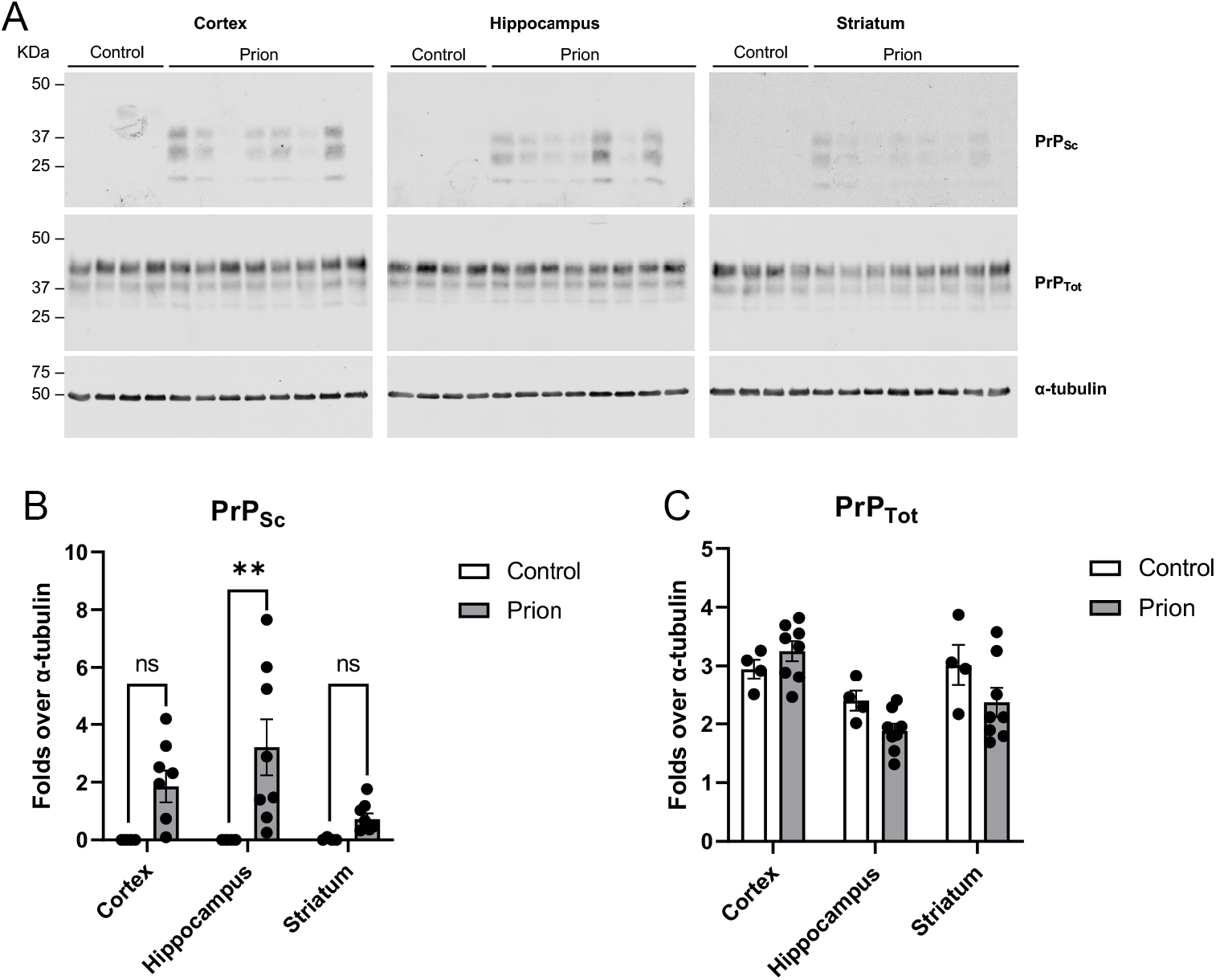
Prion-diseased mice display accumulation of scrapie prion at 7 weeks post inoculation (w.p.i.). **(A)** Lysates from cortex, hippocampus and striatum of control or prion-infected mice 7 w.p.i. were incubated in the presence or absence of proteinase K prior to western blot to detect non-digested scrapie prion protein (PrP_Sc_) and total prion protein (PrP_Tot_), respectively. Band intensity quantification for PrP_sc_ (**B**) and PrP_Tot_ (**C**) expressed as fold change over α-tubulin expression are shown as means ± SEM (n=4-8, individual values are displayed within bars; **P<0.01; two-way ANOVA Sidak Multiple comparisons).

**Supplementary figure S2.**
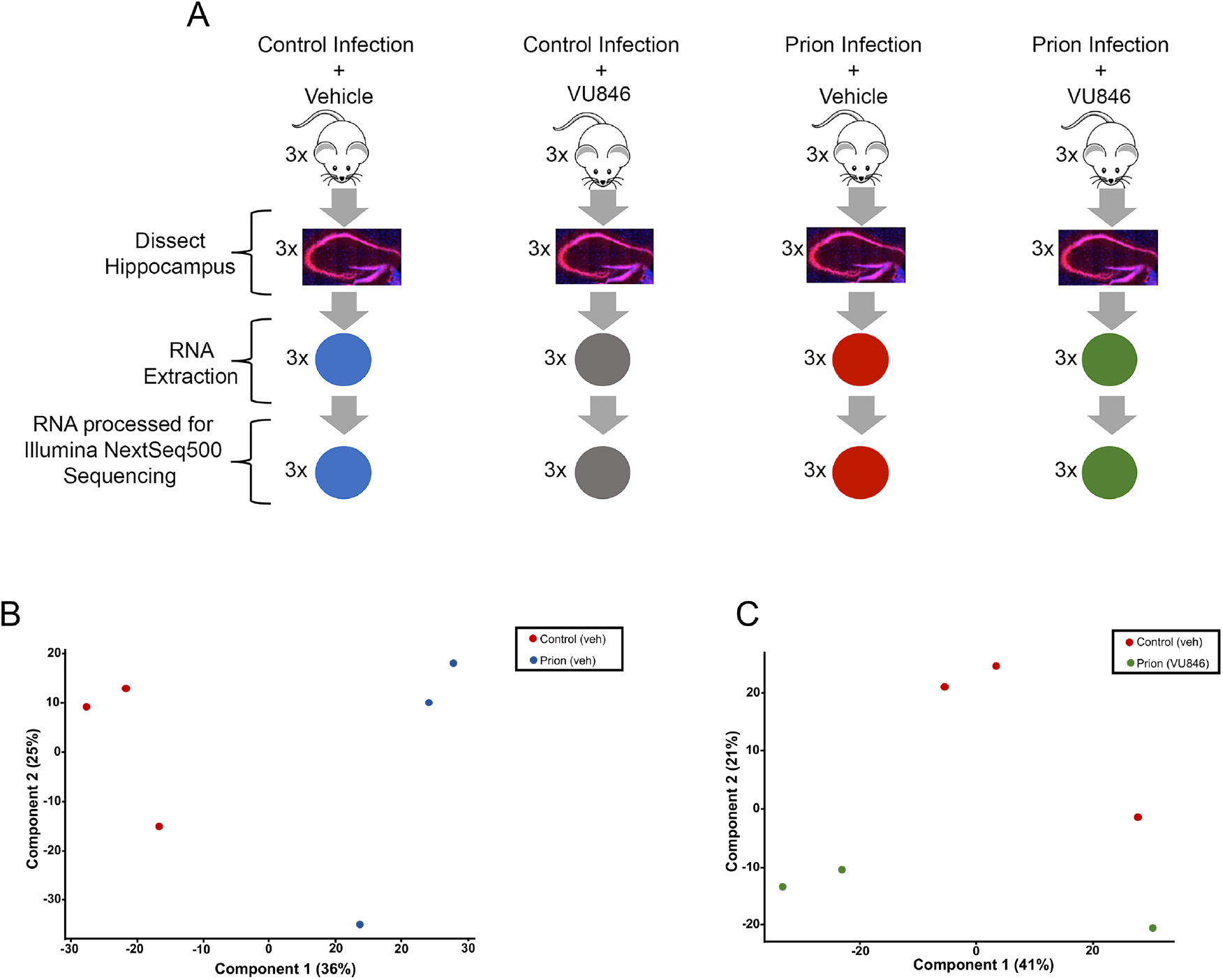
(**A**) Illustrated summary of the experimental outline and sample preparation for transcriptomics study. Hippocampi from three mice from each of four experimental groups (control + vehicle (Veh), control + VU846, prion + vehicle and prion + VU846) were dissected and the RNA extracted and processed for Illumina NextSeq500 sequencing. (**B-C**) Principal components analysis (PCA) of the transcriptomics study of **(B)** 3 control vehicle treated and 3 prion-infected animals treated vehicle and **(C)** 3 control vehicle treated and 3 prion-infected animals treated VU846.

**Supplementary figure S3.**
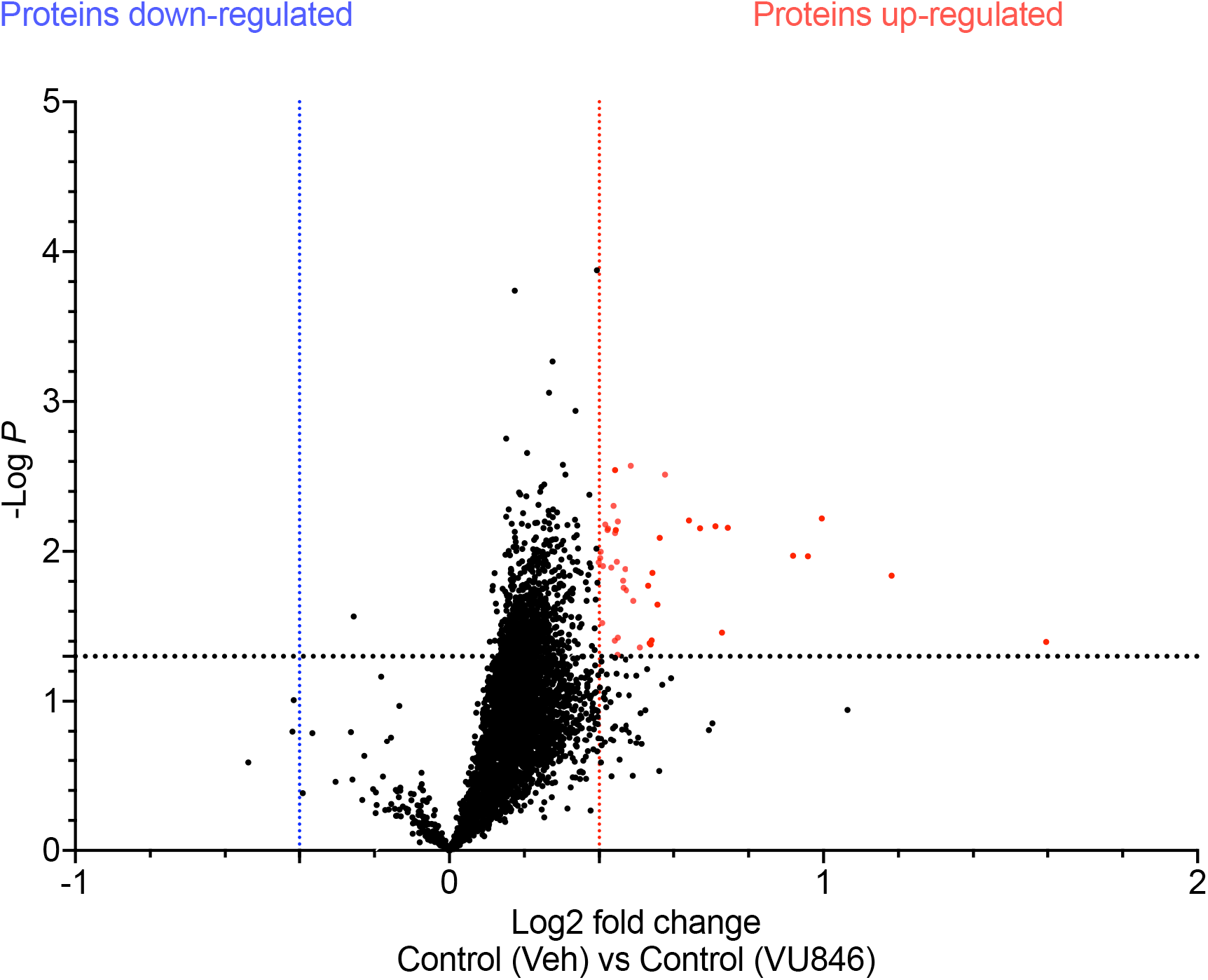
Proteomic *PAM-effect* in the control (non-infected) mice. Volcano plot representation of differential protein expression in the control-vehicle versus control-VU846. Red and blue points represent the proteins with significantly increased and decreased expression, respectively, in control-VU846 versus control-vehicle (FDR<0.05, ±Log_2_ 0.4-fold change).

**Supplementary figure S4.**
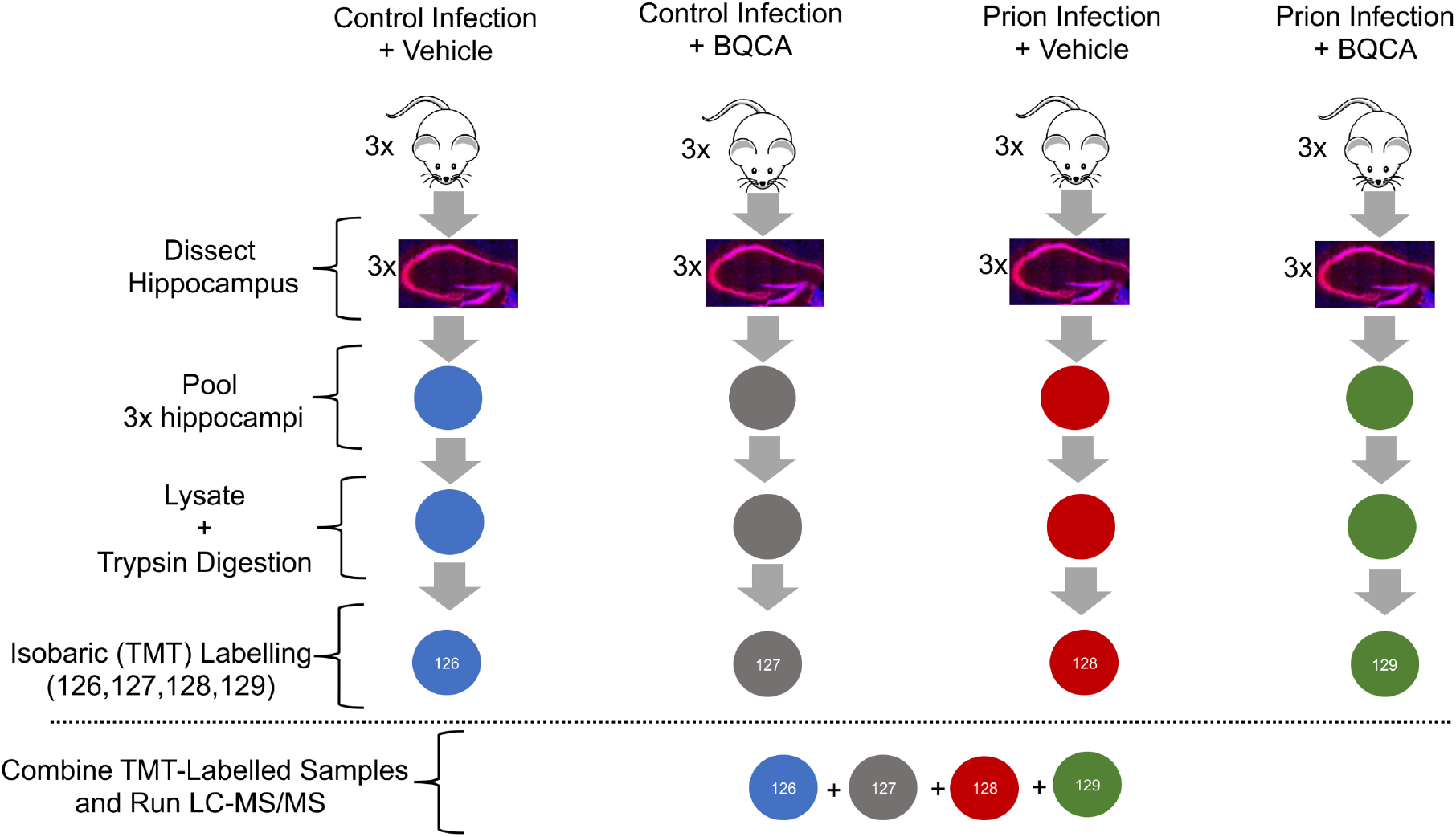
Illustrated summary of the experimental outline and sample preparation for BQCA mass spectrometry-based proteomics analysis. Hippocampi from three mice in each of the experimental groups (control + vehicle, control + VU846, prion + vehicle and prion + VU846) were pooled together, processed and labelled with respective TMT, combined and analysed by LC-MS/MS to give one experimental run.

**Supplementary figure S5.**
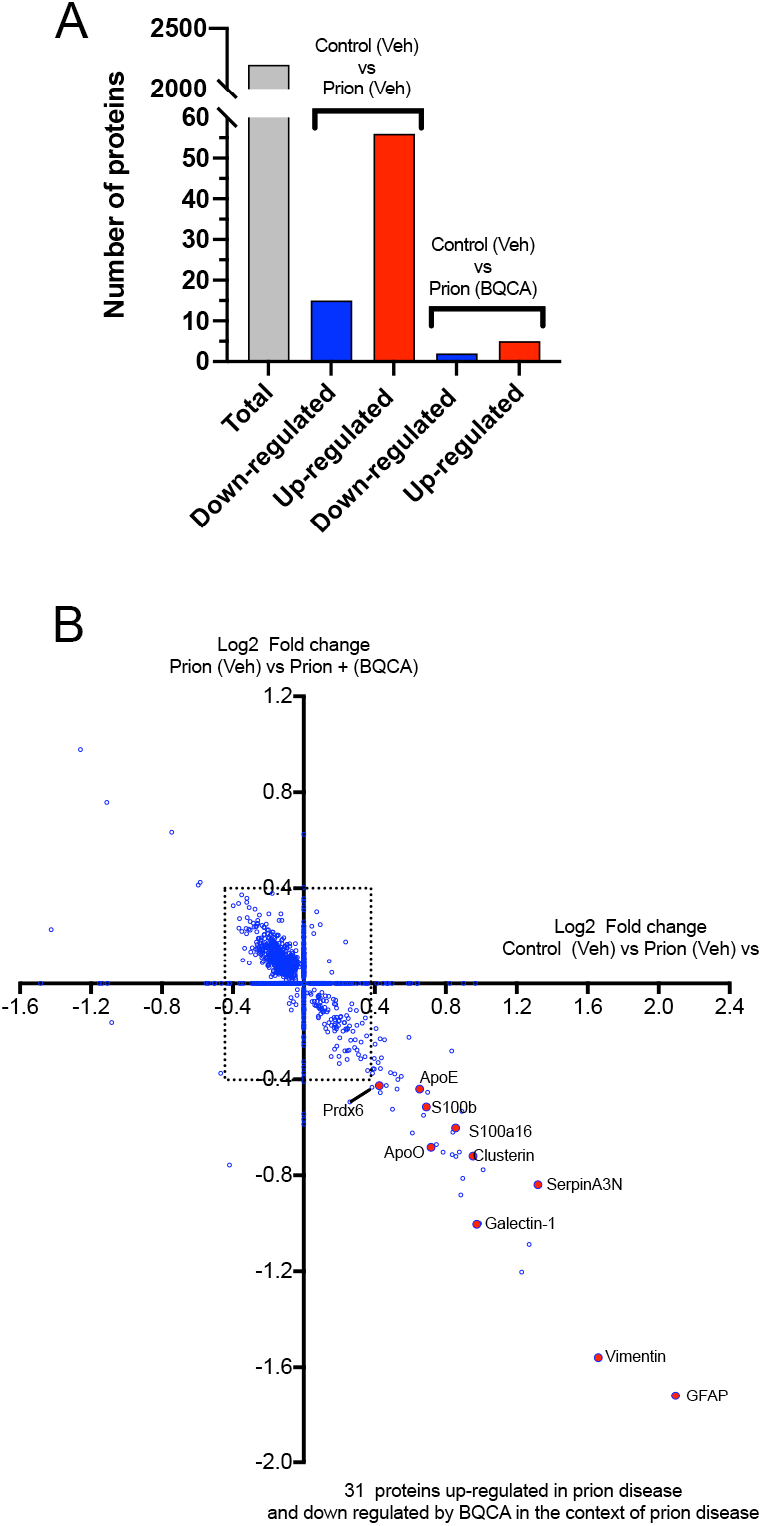
Different M_1_-PAMs mediate similar effects in prion disease. (**A**) Number of proteins that are up- and down regulated in the *prion-effect* of vehicle treated mice and mice treated with BQCA. (**B**) Quadrant scatter plot showing the effect of BQCA in the context of prion. The x-axis shows fold changes of proteins that are up- or down-regulated by prion disease (comparison between control-vehicle and prion-vehicle; *prion-effect*) and the y-axis depicts fold changes of proteins that are up- or down-regulated by BQCA in prion disease (comparison between prion-vehicle and prion-BQCA; *PAM-effect*). Proteins outside the square box are significantly changed in expression (FDR<0.05, ±Log_2_ 0.4-fold change).

**Supplementary figure S6.**
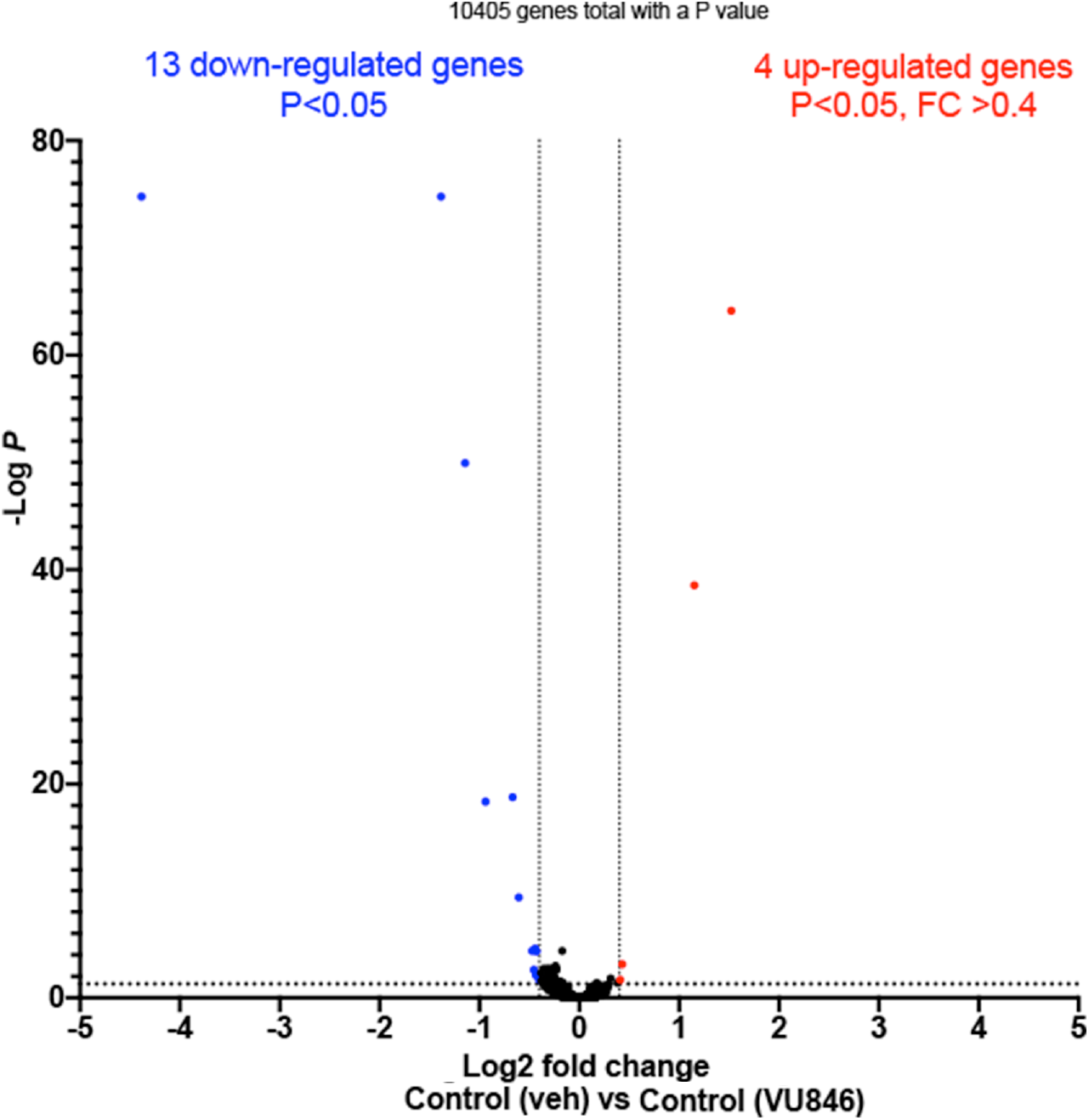
Transcriptomic *PAM-effect* in the control (non-infected) mice. Volcano plot generated using DESeq2 differential gene expression analysis of M_1_-PAM in the context of control (non-infected mice). Red and blue points represent genes with significantly increased and decreased expression, respectively, in control-vehicle versus control-VU846 (FDR<0.05, ±Log_2_ 0.4-fold change).

